# Hyperglycemia selectively increases cerebral non-oxidative glucose consumption without affecting blood flow

**DOI:** 10.1101/2024.09.05.611035

**Authors:** Tyler Blazey, John J. Lee, Abraham Z. Snyder, Manu S. Goyal, Tamara Hershey, Ana Maria Arbeláez, Marcus E. Raichle

**Author notes:** <u>**Corresponding author**</u>, Marcus Raichle, Mallinckrodt Institute of Radiology, Washington University School of Medicine, 4525 Scott Avenue, Campus Box 8131, St. Louis, MO 63110 USA.

## Abstract

Multiple studies have shown that hyperglycemia increases the cerebral metabolic rate of glucose (CMRglc) in subcortical white matter. This observation remains unexplained. Using positron emission tomography (PET) and euinsulinaemic glucose clamps, we found, for the first time, that acute hyperglycemia increases non-oxidative CMRglc (i.e., aerobic glycolysis (AG)) in subcortical white mater as well as in medial temporal lobe structures, cerebellum and brainstem, all areas with low euglycemic CMRglc. Surprisingly, hyperglycemia did not change regional cerebral blood flow (CBF), the cerebral metabolic rate of oxygen (CMRO_2_), or the blood-oxygen-level-dependent (BOLD) response. Regional gene expression data reveal that brain regions where CMRglc increased have greater expression of hexokinase 2 (*HK2*). Simulations of glucose transport revealed that, unlike hexokinase 1, *HK2* is not saturated at euglycemia, thus accommodating increased AG during hyperglycemia.

## Introduction

The brain receives glucose through facilitated diffusion across the blood-brain barrier. As a result, hyperglycemia increases the concentration of glucose in the brain as well as in the blood (1,2). Despite the increase in glucose availability, the traditional view is that hyperglycemia does not increase the cerebral metabolic rate of glucose (CMRglc) because hexokinase 1 (*HK1*), the enzyme responsible for catalyzing the first step in glycolysis, is saturated at typical blood glucose concentrations (1,3). However, positron emission tomography (PET) studies in acutely hyperglycemic normal adult humans have consistently revealed two main findings: 1) quantitative estimates of white matter CMRglc increase by approximately 40 to 50% in humans (4,5); and 2) the gray-white contrast of whole-brain normalized CMRglc images decreases, with cortical gray matter uptake in areas generally targeted by Alzheimer’s disease (AD) decreasing relative to the rest of the brain (6–8). Although these findings appear to be contradictory, a simple explanation is that hyperglycemia decreases cortical FDG uptake in whole-brain normalized PET images because whole-brain CMRglc increases slightly due to increased white matter CMRglc. This is consistent with a previous report of increased whole-brain CMRglc during acute hyperglycemia (4).

Hyperglycemia provoked by insulin resistance is a defining feature of type-2 diabetes mellitus (T2DM). Hyperglycemia is also induced by stress (e.g., see (9)). T2DM substantially increases the risk of Alzheimer’s disease (AD) (10) and stress significantly accelerates the progression of AD in amyloid precursor protein transgenic (*Tg*2576) mice (11). An element of the risk for AD imposed by hyperglycemia is that hyperglycemia promotes amyloid beta (A_β_) release in the brain by stimulating neuronal activity (12). All these observations have motivated our research which focuses on understanding how acute hyperglycemia regionally affects brain metabolism, circulation and related cellular activity in normal adult humans. We deem this a necessary component of data needed to understand how T2DM and stress contribute to the risk for and progression of AD.

To determine whether acute hyperglycemia increases fully oxidative CMRglc or aerobic glycolysis (AG), we measured both regional CMRglc and CMRO_2_ during euglycemic and hyperglycemic glucose clamps. Blood insulin levels were maintained at euglycemic levels during both clamps by infusing the somatostatin analog octreotide. Because the release of A_β_ by hyperglycemia has been attributed to an increase in neuronal activity (12) we also measured cerebral blood flow (CBF) and task-evoked changes in the blood-oxygen-level-dependent (BOLD) response (13). We then combined regional gene expression data from previously datasets (14,15), with a model of glucose transport (3) to provide a mechanistic explanation of our results.

## Methods

### Participants

Twenty-nine healthy young adults of both sexes were recruited through the Volunteers for Health program at Washington University in Saint Louis. All participants were right-handed and in good health based on a medical history and physical examination. None were taking medications (aside from oral contraceptives). All had normal fasting plasma glucose, creatinine concentrations, and hematocrits. None of the individuals had a personal history or first-degree relative with diabetes, or a personal history of significant psychiatric, neurological or cardiovascular conditions. Each participant gave their written informed consent to participate in this study, which was approved by the Radioactive Drug Research Committee and the Institutional Review Board of the Human Research Protection Office at Washington University in accord with the Helsinki Declaration of 1975.

### Experimental design

We conducted a two-arm experimental study, where subjects underwent PET-MRI to assess regional brain glucose metabolism, oxygen consumption, and blood flow during a euglycemic (90-100 mg·dL^-1^) or hyperglycemic (250-300 mg·dL^-1^) glucose clamp. Glucose was maintained at the desired level levels by 1) suppressing endogenous insulin and glucagon secretion with the infusion of octreotide, a somatostatin analog (16) and 2) using a variable-rate infusion of 20% dextrose to bring plasma glucose to the desired concentration. Subjects remained supine throughout the entire study with eyes closed during the scans. Additional details on the clamping procedure can be found in the **Supplemental Methods**.

### Image acquisition

PET and MRI data were acquired simultaneously using a Siemens Biograph PET/MRI. Cerebral oxygen metabolism and blood volume were assessed with the inhalation of approximately 25 mCi of [^15^O]O_2_ or [^15^O]CO respectively. Next, approximately 25 mCi of [^15^O]H_2_O was injected for the measurement of cerebral blood flow. PET scans were obtained twice for all three tracers, if possible. All [^15^O] tracer imaging data was acquired in dynamic listmode starting prior to tracer administration. [^15^O]H_2_O and [^15^O]O_2_ scans lasted for five minutes, whereas the [^15^O]CO scan lasted for seven minutes. Following the [^15^O] scans, 5 mCi of [^18^F]FDG was injected, and cerebral glucose metabolism was measured using a 60-minute dynamic PET scan. PET scans were obtained in 3D mode.

Functional MRI (fMRI) scans were acquired prior to the PET data using a 2D echo-planar imaging (EPI) sequence with 154 volumes (TR = 2.0 s). Imaging parameters were as follows: 64 × 64 acquisition matrix, 32 slices, 4 mm isotropic voxels, a TE of 27 milliseconds, and a flip angle of 90 degrees. Concurrent with the [^15^O]H_2_O PET scans, quantitative CBF was measured using pseudo-continuous arterial spin labeling (pCASL) (17). 2D pCASL acquisition included 80 volumes, each with 20 slices, voxel size 3.4 × 3.4 × 5.0 mm, and a 64 × 64 mm in-plane acquisition matrix. The TR was 4.0 seconds, the TE 12.0 milliseconds, and the GRAPPA acceleration factor was 2.0. Two separate pCASL runs were acquired during each study visit.

### Task Design

fMRI was used to assess changes in brain activity during a face-name encoding task prior to collecting PET data. During the first two runs, participants were shown a picture of a human face with a name underneath it and asked to indicate via button press if the name matched the face. The same combination of face and names was used for both runs. For the final two runs, participants were presented with faces without the names underneath and asked if they remembered the name that was associated with that face during the first two runs. A different set of names/faces was used for the hyperglycemic visit than was used for the euglycemic visit. List assignment was counterbalanced across participants.

### Image Processing

Additional details on image processing can be found in the **Supplemental Methods**.

### MRI

Initial preprocessing of the pCASL data included odd-even slice intensity correction (18), motion correction (19), registration to MNI152 2mm atlas space with FSL (20), and high pass temporal filtering (0.08 Hz). Quantitative CBF images were then created using sing subtraction (21) and a one compartment model (22).

All processing of the task fMRI data was conducted using FEAT in FSL 6.0.0 (20). Preprocessing included motion correction (19), brain extraction, spatial smoothing with a 5-mm full-width half-maximum Gaussian kernel, global normalization of the entire 4D timeseries, high-pass temporal filtering with a cutoff of 100 seconds, and alignment to MNI152 space. Voxelwise group analyses were performed by randomly permutating the glycemic condition within a subject 10,000 times (23). Multiple comparisons correction was performed using threshold-free cluster enhancement (24). ROIs were constructed by placing a 10 mm sphere on 4 task-activated regions (**Figure S9**) and 4 task-deactivated regions (**Figure S10**).

#### PET

The NiftyPET software platform was used to reconstruct the dynamic PET data (25). Reconstruction included correction for scatter, attenuation, and motion. Each subject’s PET images were then resampled to MNI152 2mm atlas space using a combination of in-house tools (26) and ANTS (27). Standardized uptake value ratios (SUVRs) were used to assess regional changes in metabolism. SUVR is a semi-quantitative measure where the tracer uptake in each brain region is reported relative to a reference region. Although our laboratory traditionally uses a whole-brain reference region (28), the global increase in glucose consumption (**Table S3**) prevented us from doing this. Instead, we chose the three FreeSurfer (29) defined regions (middle temporal, lateral orbitofrontal, postcentral gyrus) with the smallest absolute change in CMRglc (**Figure S2A-B**). Using this empirically defined reference region greatly increased the similarity between our quantitative and SUVR group average difference (hyperglycemia – euglycemia) images (**Figure S2C-E**). Prior to any comparisons, atlas space SUVR images were smoothed with a 5 mm FWHM gaussian kernel.

Kinetic modeling was also used to quantify CMRglc in a subset of participants (n=16) in whom very fast hand drawn arterial samples were acquired during the [^18^F]FDG scan. At the whole brain level, we used non-linear least squares to fit a reversible two-compartment model the dynamic [^18^F]FDG data (30). The model includes four rates constants (K_1_, k_2_, k_3_, and k_4_) and an additional term, V_b_, that accounts for arterial blood volume. K_1_ and k_3_ govern influx from the blood and first compartment, respectively. Loss of tracer from the first compartment is governed by k_2_, and efflux from the second compartment to the first is designated by k_4_. To fit this model at the voxel level, we used the hierarchical-basis function method (31), with the additional of a spatial constraint. Following the recommendation of Wu et al. (32), a lumped constant (LC) of 0.81 was used to calculate CMRglc.

#### Gene expression

Regional microarray expression data were obtained from 6 post-mortem brains (provided by the Allen Human Brain Atlas (https://human.brain-map.org; (33)). Data were processed with the abagen toolbox (version 0.1.3; https://github.com/rmarkello/abagen) using the 48-region volumetric atlas in MNI space used for PET imaging.

Cell-type specific analyses were performed using two independent datasets containing single-cell RNA expression from individual brain cells (14,15). The first dataset, which was described by Siletti et al., consists of 3,369,219 cells taken from 10 different brain regions (14) Although most of the dissections targeted neuron rich regions, samples contained cells from both white and gray matter. The Siletti et al. data was from https://github.com/linnarsson-lab/adult-human-brain. The second dataset was originally published by Seeker et al. (15), and was downloaded from https://cellxgene.cziscience.com/collections/9d63fcf1-5ca0-4006-8d8f-872f3327dbe9. The Seeker et al. data consisted of 45,528 cells taken specifically from white matter in four separate brain regions. For both datasets, classification of cell class (neuronal vs. non-neuronal) and cell type (astrocytes, microglia, etc.) was based on expression clusters defined in the original reports. The expression matrices (Genes × Cells) were binarized and the number of cells expressing each hexokinase isoform (*HK1*, *HK2*, *HK3*, and *GCK4*) was then quantified for each cell type over all available regions. For a full description of the datasets see the original reports (14,15).

### Glucose transport modeling

Glucose transport was modeled using the carrier model introduced by Barros et al. (3). The basic unit of the model is a glucose carrier that can face either side of a membrane and exist in either a bound or unbound state. This model leads to four differential equations with four compartments and eight rate constants per carrier (**Figure S12**). The number of rate constants is reduced by half by assuming the rate constants are symmetric across the membrane. The first two rate constants, f_1_ and f_2_, describe the transport of the unbound and bound carrier across the membrane. The remaining constants describe the association (K_on_) and dissociation (K_off_) of glucose from. Following Barros et al. (3), we used the following values for each rate constant: f_1_ = 200 s^-1^, f_2_ = 3000 s^-1^, K_on_ = 1E8 M^-1^s^-1^, K_off_ =4E6 s^-1^.

The full model consists of three separate carriers, with a carrier separating each of the following membranes: 1) blood : endothelium, 2) endothelium : interstice, and 3) interstice : intracellular. The same rate constants were used for each carrier. The concentration of the three carriers was fixed at a 1:1:10 ratio (3). Once inside the intracellular compartment breakdown of glucose was modeled using Michaelis-Menten kinetics:

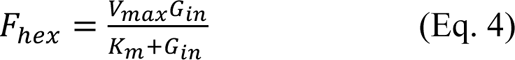

where G_in_ is the intracellular concentration of glucose. Gray matter simulations used a V_max_ of 6.8 μM s^-1^, a K_m_ of 50 mM, and a total carrier concentration of 2 μM in the interstice : intraceullar membrane (3). During euglycemia, these parameter values resulted in a G_in_ of 1.2 mM and a F_hex_ of 6.5 μM s^-1^. For white matter simulations, we varied both K_m_ and the G_in_ at euglycemia to determine the model’s sensitivity to each parameter (**Figure 7A**). The values for V_max_ and carrier concentration were set to ensure that at euglycemia G_in_ matched the chosen value and F_hex_ was ∼1.7 smaller in white matter than in gray matter, mirroring the difference in CMRglc between white and gray matter in our study.

### Statistics

Unless otherwise stated, statistical analyses were done using the R programming language (34). A linear mixed model with a fixed effect for clamp condition and a random intercept for subject was used to determine the statistical significance of differences in brain metabolism between conditions. No other covariates were added to the model. The voxelwise mixed models were fit using the nlme package (35). The difference between conditions was considered significant when the *p*-value was less than 0.05. Multiple comparison across space were accounted for by controlling the False Discovery Rate at 0.05.

The nlme package was also used to perform a piecewise linear regression of the plasma glucose and insulin data that were obtained during the glucose clamps. The fixed effects included time, clamp condition, and the interaction between time and condition. There were two regressors, one for each condition, that allowed for the slope of the regression line to vary after a 55 minutes from the start of the glucose clamp. We also included a random intercept for study visit nested within subject. Slopes that were different from 0 at the *p* < 0.05 level (no correction for multiple comparisons) were considered significant. Pearson correlation was used to quantify the degree of spatial correspondence between baseline metabolism and the change induced by hyperglycemia. All reported values are means and 95% confidence intervals (CIs) unless otherwise stated.

## Results

### Participants

Our dataset consists of twenty-nine healthy young adults (15 female, 14 male), mean age of 34.5 + 10.1 years, and a mean body mass index (BMI) of 25.3 + 3.3 kg/m2. Most, but not all (19/29), participants had imaging data for both euglycemia and hyperglycemia conditions. The median time between experimental visits (euglycemia or hyperglycemia) was 49 days with a range of 12 – 966 days. The total number of participants for each metabolic parameter of interest is listed in **Table S1**.

### Blood glucose and insulin levels

The target plasma glucose concentration was 90-100 mg·dL^-1^ for the euglycemic clamp and 250- 300 mg·dL^-1^ for the hyperglycemic clamp. A piece-wise linear mixed model was used to fit the plasma glucose and insulin data obtained during each condition (**Figure 1**). Baseline plasma glucose (83.6 ± 8.0 mg·dL^-1^) and insulin (43.8 ± 20.1 pmol·L^-1^) did not differ across conditions (*p* > 0.67). After 55 minutes, the average plasma glucose level was near the target concentration for both the hyperglycemic (301.5 ± 11.29 mg·dL^-1^) and the euglycemic (104.3 ± 5.5 mg·dL^-^1) clamp (**Figure 1A**). However, in some individuals, plasma glucose rose slightly above the target concentration during the beginning of the glucose clamp and then slowly decreased throughout the scan (**Figure 1A**). On average, plasma insulin levels increased slightly during the hyperglycemic clamp. Importantly, insulin levels were never greater than values typically seen after a carbohydrate meal (∼425 pmol·L^-1^) and were largely below typical fasting insulin levels (∼140 pmol·L^-1^; **Figure 1B**). These observations are quantified in **Table S2**.

**Figure 1:**
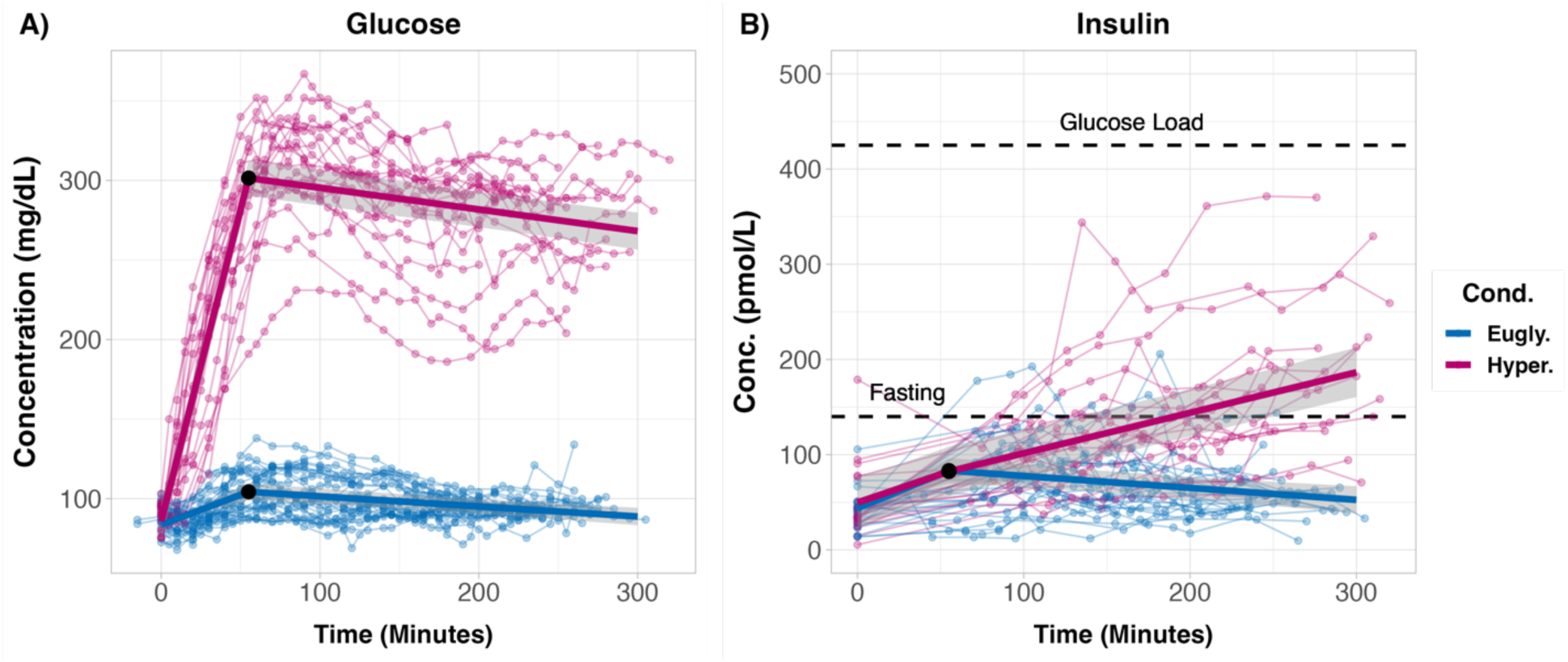
Time course of plasma glucose and insulin levels during glucose clamping. **A)** After the hyperglycemic clamp (red) the average plasma glucose level was approximately 300 mg·dL^-1^, whereas it remained near 100 mg·dL^-1^ during the euglycemic clamp (blue). A piecewise linear regression with a breakpoint at 55 minutes (black dot) was used to compute population estimates (thick lines) and their 95% CIs (gray ribbons). In both groups, the average blood glucose level increased prior to the breakpoint and then decreased afterwards (see Results). **B)** Average plasma insulin also increased in both groups prior to the breakpoint. However, after the breakpoint insulin decreased during the euglycemic clamp and increased in the hyperglycemic clamp. The top dashed black line indicates a published value for the peak plasma insulin concentration after a 75 gram oral glucose tolerance test (92). Note that even though insulin increased throughout the hyperglycemic clamp, it always remained below the peak value form the oral glucose tolerance test. For comparison, the lower dashed black line is the 97.5% reference limit for fasting insulin (93). The light lines and dots are data from individual sessions.

### Global and regional cerebral glucose consumption

To determine if glucose consumption is altered by acute hyperglycemia, we first obtained quantitative estimates of glucose metabolism for the brain as a whole (**Table S3**). Hyperglycemia significantly increased whole brain CMRglc (increase of 2.84 ± 2.41; *p* = 0.042), raising it from 30.70 ± 1.90 μMol·hg^-1^·min^-1^ at euglycemia to 33.54 ± 1.88 μMol·hg^-1^·min^-1^. Hyperglycemia also altered the kinetics of [^18^F]FDG (**Table S3**). All four of the kinetic model rate constants significantly differed between conditions (*p* < 0.001). In particular, K_1,_ which describes the transport of [^18^F]FDG from the blood to the brain, was much lower during hyperglycemia (6.61 ± 1.00 mL·hg^-1^·min^-1^) than during euglycemia (12.67 ± 1.01 μMol·hg^-1^·min^-1^).

We also computed voxelwise CMRglc to examine regional differences in the brain’s response to acute hyperglycemia (**Figure 2A)**. As expected, the effect of hyperglycemia on CMRglc was regionally specific. Significant elevations in CMRglc were largely restricted to white matter, brain stem, the cerebellum, and the medial temporal lobe (*p* < 0.05; False Discovery Rate corrected) (**Figure 2B)**. Regional increases in CMRglc were especially robust in deep white matter; this finding was present in every subject in which the comparison was possible (**Figure 2C**). Further analysis revealed a tight inverse relationship between regional change in CMRglc and baseline glucose consumption (r = 0.94). Thus, regions with low basal metabolic rate exhibited the greatest increases in CMRglc (**Figure 2D).** CMRglc was unchanged throughout most of cortical and subcortical gray matter. Small clusters in both cortical (e.g., precuneus) and subcortical (e.g., putamen) gray matter showed CMRglc decreases (**Figure 2B).** Nearly all these decreases, but not the increases, in CMRglc were eliminated by adjusting CMRglc for potential changes in the lumped constant (a correction factor that accounts for differences in the transport between [^18^F]FDG and glucose) during hyperglycemia (see **Supplemental Methods**; **Figure S1**).

**Figure 2:**
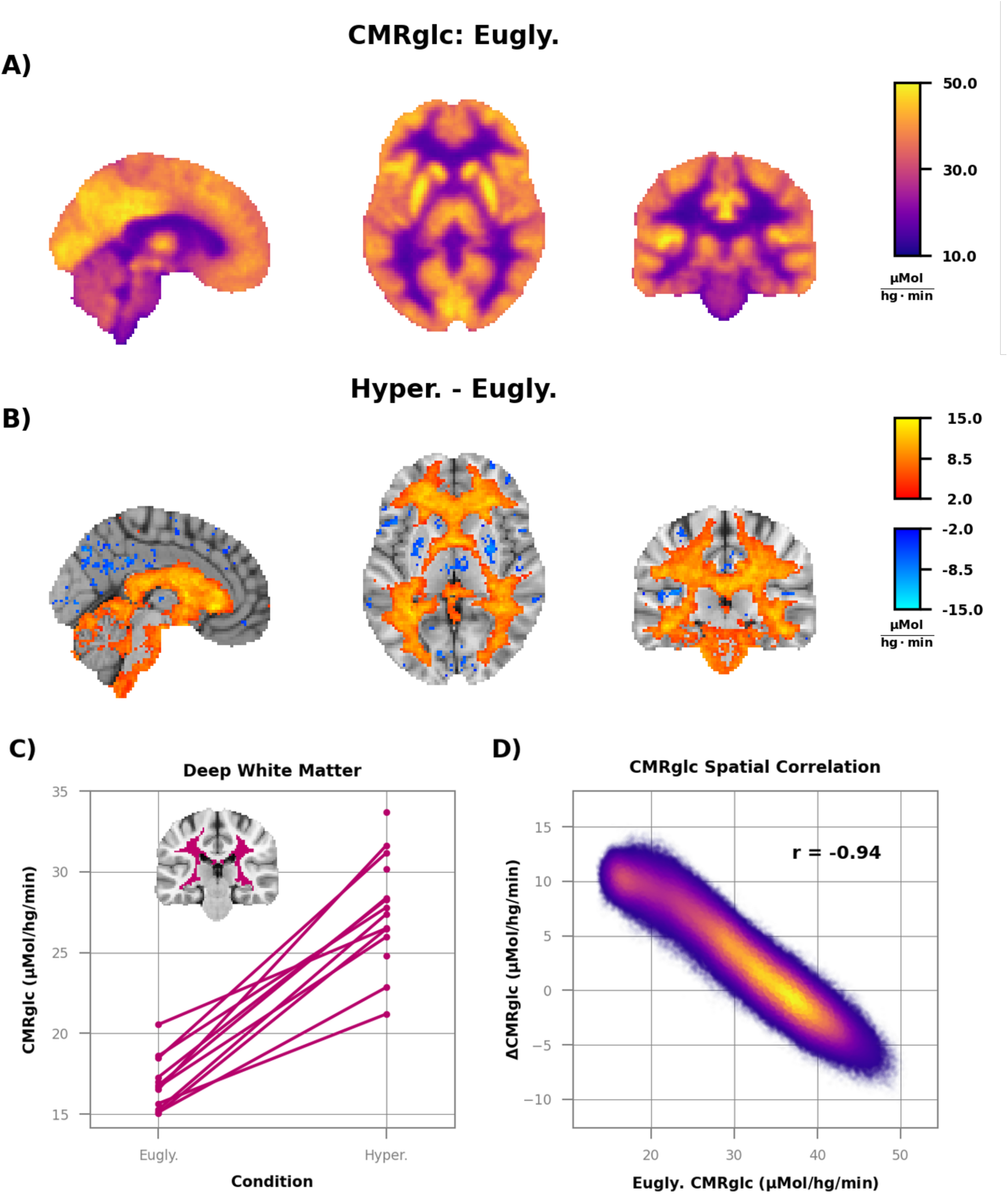
Quantitative increases in CMRglc in regions with low basal metabolic rates. **A)** Group average image of CMRglc during euglycemia (n=13) and **B)** difference between hyperglycemia (n=14) and euglycemia. Only voxels where the difference is different from zero (*p* < 0.05; False Discovery Rate corrected) are shown in color in **B). C)** CMRglc data from the deep white matter ROI (inset), showing a robust increase in glucose consumption with hyperglycemia (10.6 ± 1.6 μMol·hg^-1^·min^-1^; *p* < 0.0001). **D)** Scatterplot of CMRglc during euglycemia vs. the difference between hyperglycemia and euglycemia. Each point is a voxel and color indicates density. Note the strong negative correlation between baseline CMRglc and hyperglycemia induced change, with the regions with the smallest baseline values showing the greatest change (r = 0.94).

### Regional non-oxidative glucose consumption

We next examined regional measurements of oxygen consumption to determine if hyperglycemia increases glucose consumption through oxidative or non-oxidative pathways. Technical difficulties in obtaining [^15^O]O_2_ PET data with simultaneous arterial blood sampling precluded reporting of quantitative measurements of CMRO_2_. However, we can report SUVRs, which express regional metabolism relative to a reference region. We created a metabolically stable reference region by combining the three brain regions (middle temporal, lateral orbitofrontal, and postcentral gyrus) with the smallest absolute change in CMRglc (Methods; **Figure S2**).

We did not find any regional differences in oxygen consumption (**Figures 3)**; the group average SUVR images of hyperglycemia (**Figure 3A**) and euglycemia (**Figure 3B**) were essentially identical. Oxygen metabolism was stable even in deep white matter (**Figures 3C** where there was a robust increase in CMRglc (**Figure 2C).** There was also no spatial correlation between baseline oxygen consumption and change during hyperglycemia (**Figure 3D**), a negative finding that stands in stark contrast to the CMRglc results (**Figure 2D**). There was also no change in the regional oxygen extraction fraction (OEF) (**Figure S3**).

**Figure 3:**
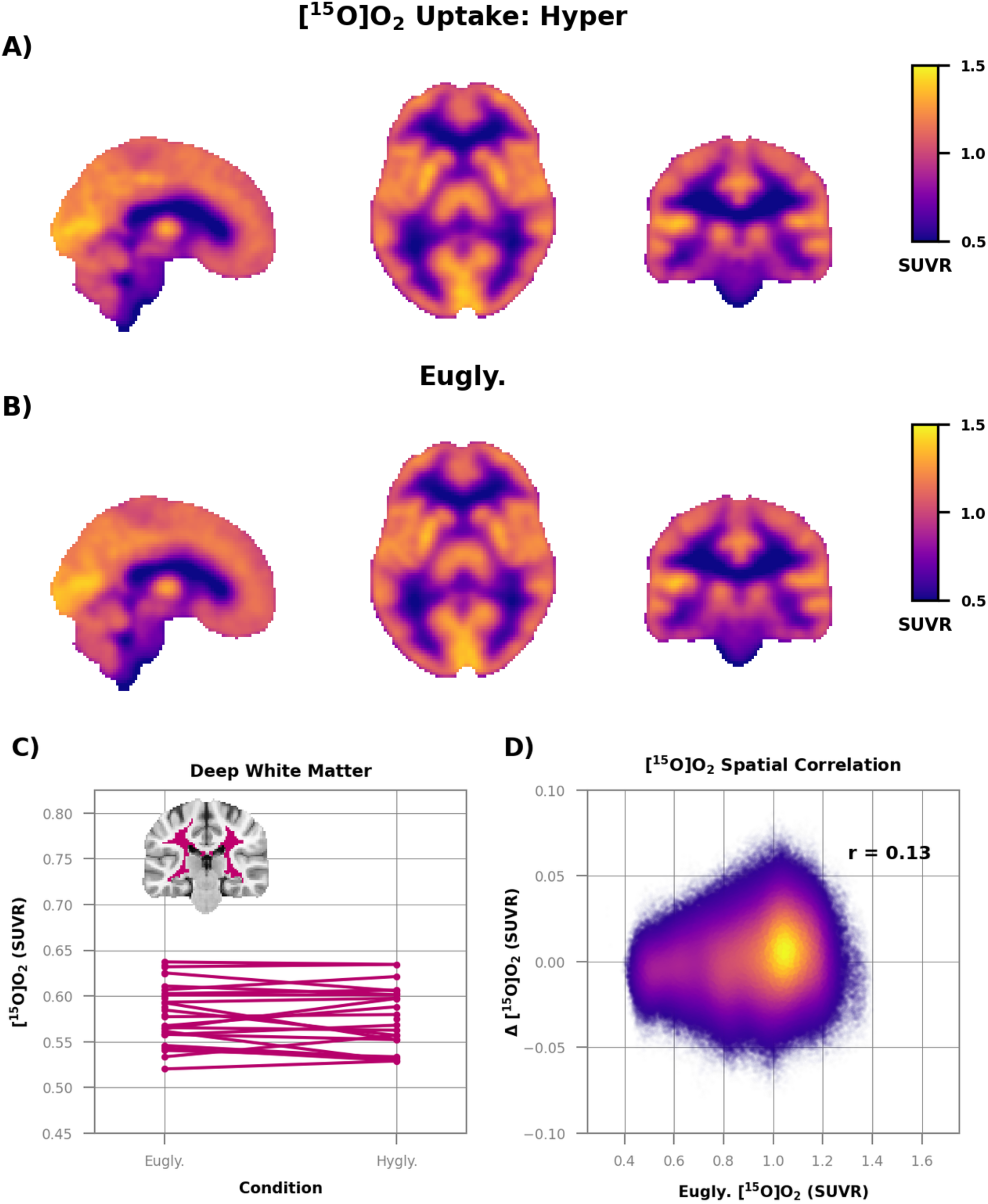
No change in relative oxygen consumption during hyperglycemia. The group average images of relative oxygen metabolism during **A)** hyperglycemia (n=21) and **B)** euglycemia (n=26) are essentially identical. No differences (*p* < 0.05; False Discovery Rate corrected) were found between the two conditions. **C)** Relative oxygen metabolism in the deep white matter. The average difference between conditions (-0.002 ± 0.007 SUVR) was not different from zero *(p* = 0.53). **D)** Scatterplot of relative oxygen consumption during euglycemia vs. the difference between hyperglycemia and euglycemia. Each point is a voxel and color indicates density. There was no relation between baseline oxygen metabolism and hyperglycemia-induced change (r = 0.13).

The observed increase in white matter glucose consumption without a change in oxygen consumption suggests that hyperglycemia increases non-oxidative glucose metabolism, often referred to as aerobic glycolysis (28). To confirm this impression, we computed the relative oxygen-to-glucose index (rOGI) as the ratio of relative oxygen consumption (**Figure 3)** and glucose consumption (**Figure S4**). By definition, rOGI inversely correlates with non-oxidative glucose consumption. Thus, a decrease in rOGI indicates an increase in non-oxidative glucose use relative to the reference region. Hyperglycemia markedly altered the contrast properties of the group average rOGI images (**Figure 4**). At euglycemia, rOGI was higher in white matter than in cortical gray matter (**Figure 4B**), whereas during hyperglycemia cortical gray matter had the higher rOGI (**Figure 4A**). This change in contrast was largely attributable to a significant decrease in rOGI (an increase in non-oxidative glucose consumption) throughout white matter, brain stem, cerebellum, and the medial temporal lobe (**Figure 4C**). Regions with low basal rOGI consumption showed the greatest absolute change with hyperglycemia (r = -0.58). Modest focal rOGI increases were observed throughout the brain, including parts of the striatum and the precuneus. These increases in rOGI largely corresponded to voxels in which quantitative (**Figure 2**) and SUVR (**Figure S4**) measurements of glucose consumption decreased.

**Figure 4:**
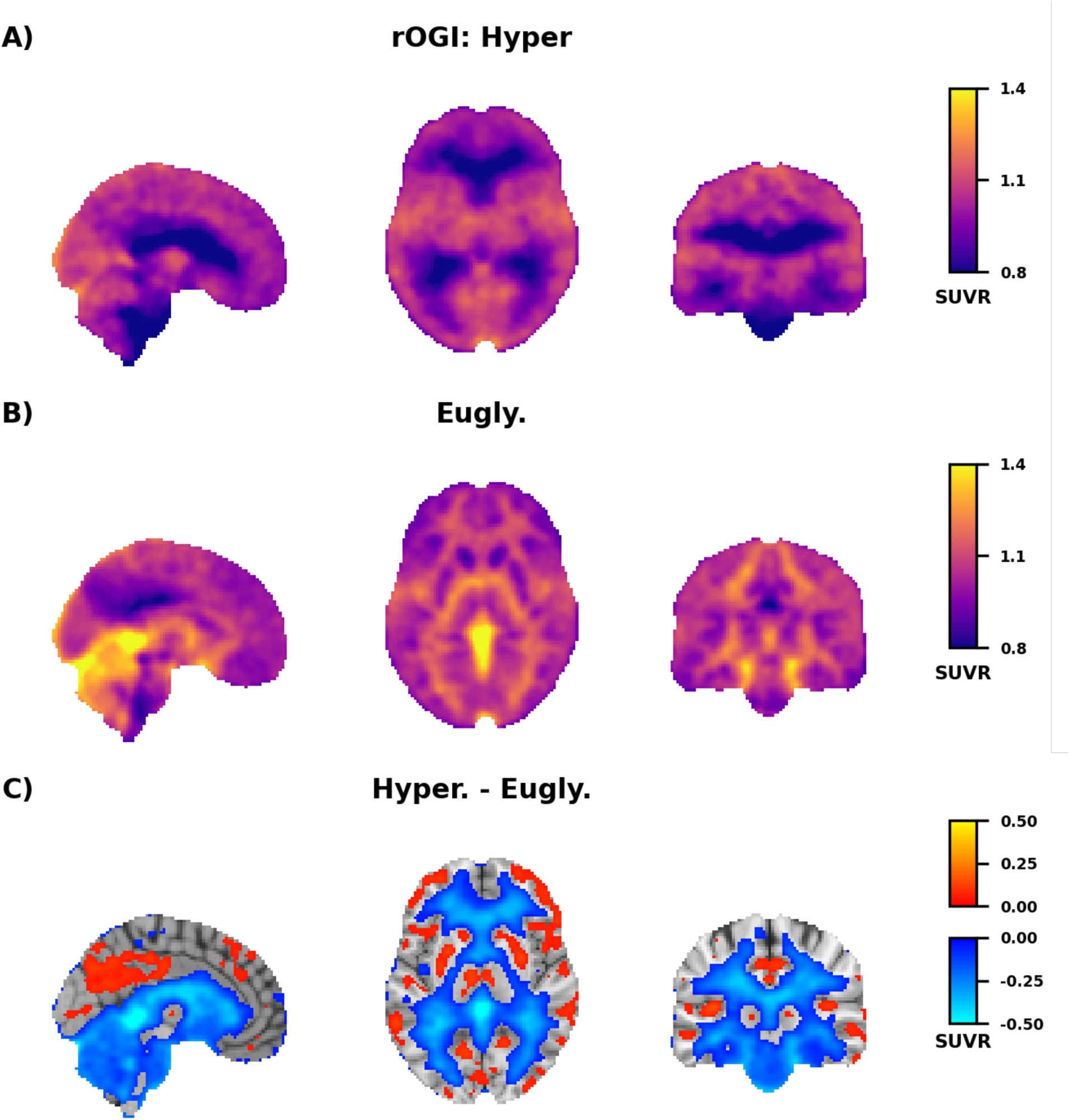
Hyperglycemia-induced changes in relative oxygen-to-glucose (rOGI) ratio. **A)** Group average rOGI during the euglycemic (n=25) and **B)** hyperglycemic (n=21) clamp. rOGI was computed by taking the ratio of the [^15^O]O_2_ SUVR and [^18^F]FDG SUVR images and then normalizing by the uptake in a reference region where CMRglc did not change during hyperglycemia (**Figure S2**). **C)** Group average difference in relative rOGI between the hyperglycemic and euglycemic clamp. Voxels different from zero after correction for multiple comparisons (*p* < 0.05; False Discovery Rate corrected) are shown in color. Voxels where rOGI decreased are shown in blue; voxels where rOGI increased are shown in orange/yellow.

### Blood flow and task-evoked activity

As neural activity usually is accompanied by an increase in non-oxidative glucose consumption (36) and cerebral blood flow (37), we next examined the possibility that acute hyperglycemia induces a change in neural activity. First, we used PET to measure CBF and blood volume (CBV). We found no changes in either CBF (**Figure 5**) or CBV (**Figure S5**) during hyperglycemia. To confirm this result, we also measured quantitative blood flow with arterial spin labeling MRI. Again, no changes were found (**Figure S6**).

**Figure 5:**
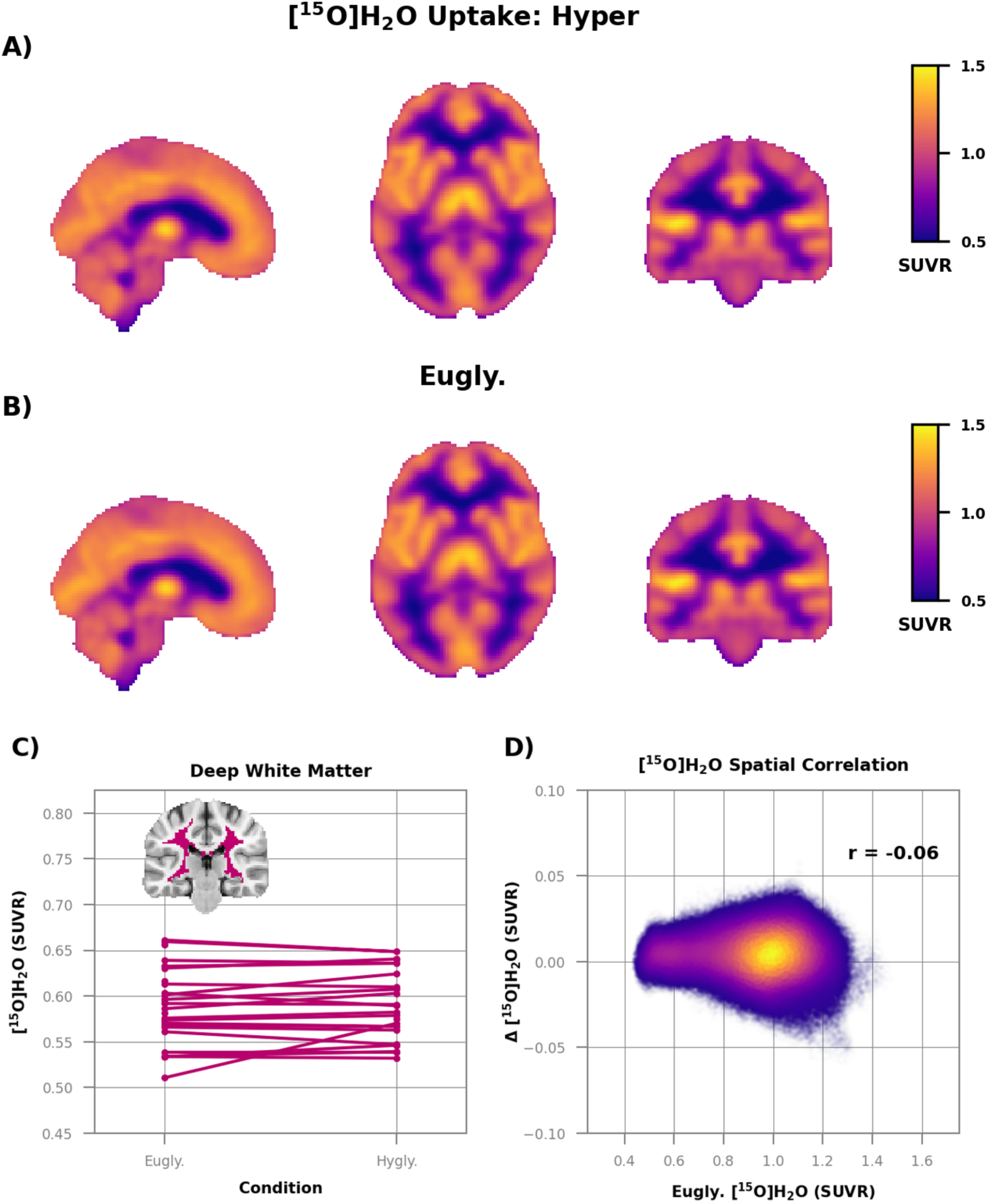
No change in relative cerebral blood flow during hyperglycemia. The group averages of relative blood flow images during A) hyperglycemia (n=21) and B) euglycemia (n=26) are essentially identical. Statistical testing showed no differences (p < 0.05; False Discovery Rate corrected) between the two conditions. Relative blood flow was computed by summing the early frames of the [^15^O]H_2_O PET images and normalizing by the uptake in a reference region where CMRglc did not change (**Figure S2**). C) Relative blood flow in the deep white matter. The average difference between conditions (0.004 ± 0.007 SUVR) was not different from zero (p = 0.24). D) Scatterplot of relative blood flow during euglycemia vs. the difference between hyperglycemia and euglycemia. Each point is a voxel and color indicates density. There was no relationship between baseline blood flow and hyperglycemia-induced change (r = -0.06).

Although we found no change in resting state blood flow, it is possible that hyperglycemia increases neuronal responses to stimuli (38). Therefore, we used fMRI to examine changes in BOLD response during a face-name task consisting of separate encoding and retrieval stages (see Methods). Although there were robust fMRI responses during both encoding (**Figure S7A**) and retrieval (**Figure S8A**) consistent with prior work (39), there were no significant hyperglycemia-associated differences either at the voxel level (**Figures S7 and S8**) or within regions of interest that were activated by the task (**Figures S9 and S10**). Task performance data revealed no significant effects of hyperglycemia (**Figure S11**).

### Regional gene expression

To investigate potential mechanism(s) underlying focal increases in glucose consumption during acute hyperglycemia, we used the Allen Human Brain Atlas (33) to examine the regional mRNA expression of hexokinase (HK), the enzyme which catalyzes the first step in glucose metabolism, and exists in several isoforms (40). *HK1*, the primary HK in the brain is highly expressed in cortical gray matter (**Figure 6A**) whereas, in white matter, *HK1* expression is low and *HK2* expression is high. Importantly, the *HK1*/*HK2* ratio was positively correlated with resting glucose consumption (r = 0.63), and negatively correlated with the change in glucose consumption during hyperglycemia (r = -0.63; **Figure 6B**).

**Figure 6:**
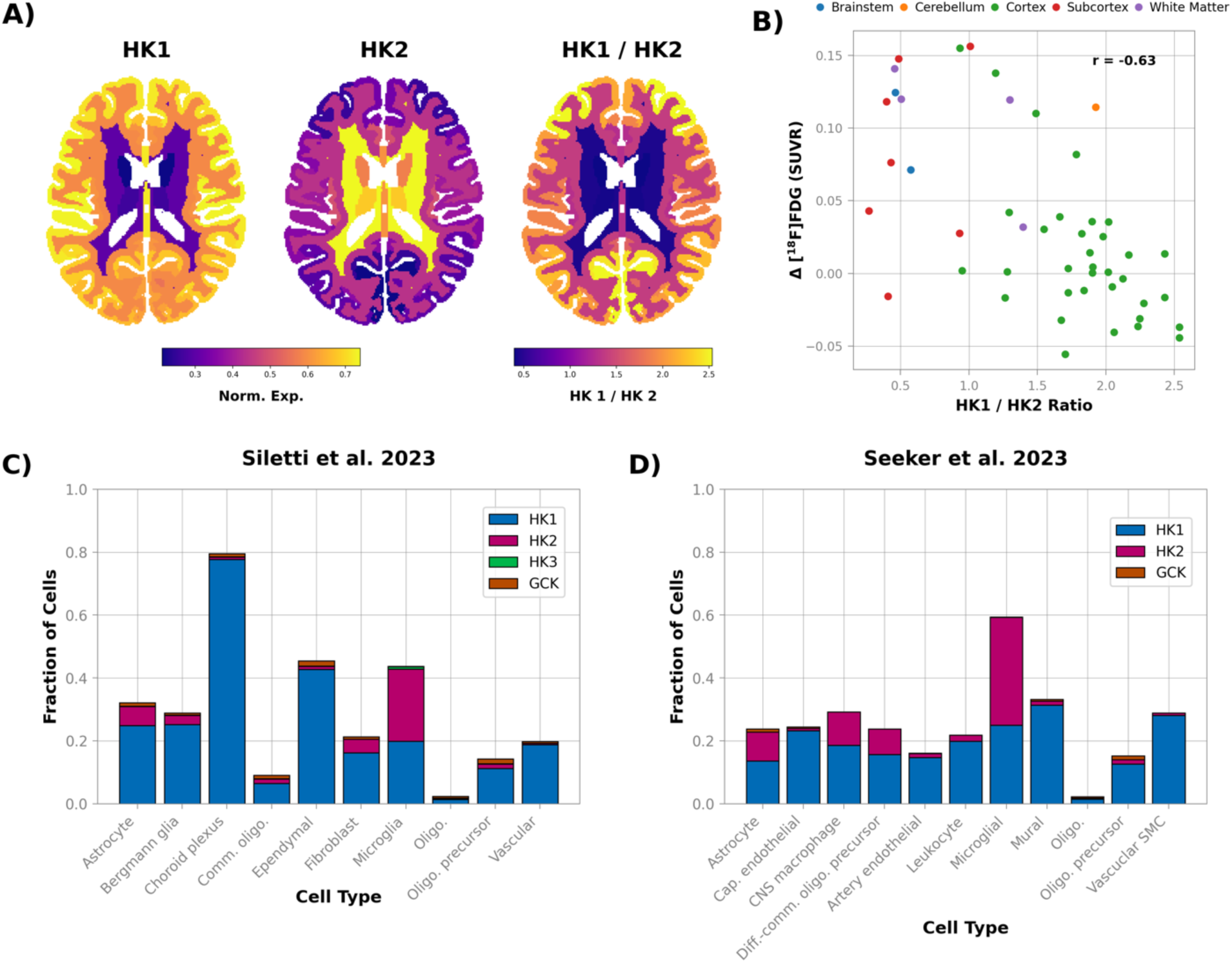
Regional gene expression and glucose consumption during hyperglycemia. **A)** Regional expression of hexokinase 1 (*HK1*) and hexokinase 2 (*HK2*) in the human brain derived from the Allen Human Brain Atlas (33). *HK1*, which has K_m_ of glucose 3-10 times lower than *HK2* (41), is expressed more in gray matter than in white matter. Conversely, *HK2* is expressed more in white matter. Note the atlas used to create the expression images was bilaterally symmetric. **B)** The *HK1*/*HK2* ratio was negatively correlated with the regional change in glucose consumption induced by hyperglycemia. **C)** and **D)** Expression of hexokinase isozymes in different cell types from two independent single-nucleus RNA expression data sets (14,15). Samples from the first study, Siletti et al. 2023, contained cells from both gray and white matter (14). The second study, Seeker et al., 2024, only contained samples from white matter (15). In both data sets *HK2* expression was the greatest in microglia, although *HK2* was also a substantial fraction of hexokinase expression in astrocytes.

Next, to determine what cell type(s) express *HK2* in the human brain, we explored two recently published single-nucleus RNA datasets (14). The first study, Siletti et al. contained samples from both white and gray matter (14), whereas the second study, Seeker et al. sampled only white matter (15). The overall fraction of cells expressing *HK2* was low in both the Siletti et al. data set (1.5%) and in the Seeker et al. data (4%). However, the cells expressing *HK2* were largely non-neuronal, with non-neuronal cells making up 64.7% of *HK2* expressing cells in the Siletti et al. data and 99.0% in the Seeker et al. data. When non-neuronal cells were grouped by cell type, we found that *HK2* expression was highest in microglia and astrocytes (**Figure 6C-D**). *HK2* expression was particularly pronounced in microglia, the only cell type in which *HK2* was expressed in more cells than *HK1*.

### Simulated glucose flux during hyperglycemia

The correlation between changes in glucose consumption and *HK2* expression suggests that differences in enzyme kinetics may play a role in regional differences in the brain’s response to hyperglycemia. To assess the likelihood of this hypothesis, we simulated flux through both gray matter (*HK1*) and white matter (*HK2*) over a range of blood glucose levels using the carrier model proposed by Barros et al. (3). This model depends on prior values for both the Michaelis constant K_m_ of *HK* for glucose and the intracellular concentration of glucose in the tissue of interest during euglycemia (see Methods). As the value for each parameter in white matter is uncertain, we first ran simulations over a range of values to determine how each parameter affects the stimulated change in flux between euglycemia and hyperglycemia. We found that during hyperglycemia, flux, defined as the irreversible phosphorylation of glucose to glucose-6- phosophate, increases as K_m_ increases, whereas flux decreases as the basal intracellular glucose concentration increases (**Figure 7A**).

**Figure 7:**
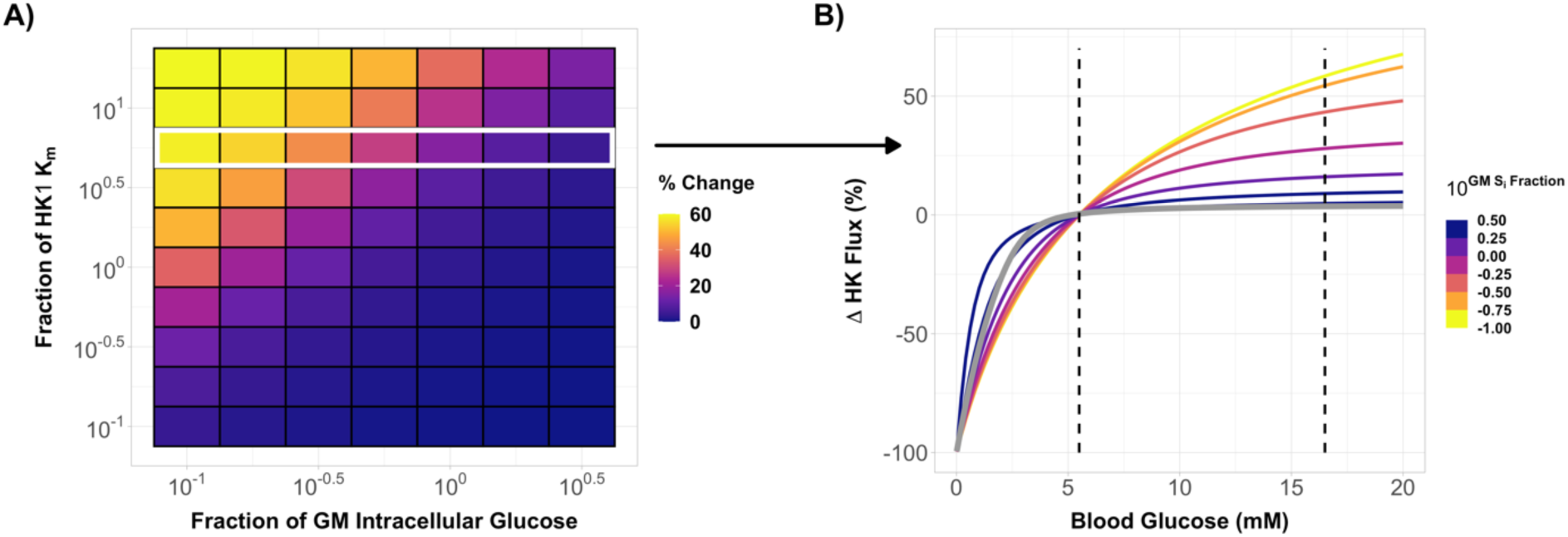
Simulations of glucose transport predict increased flux through *HK2*, but not *HK1*, during hyperglycemia. **A)** A model of glucose transport (3) was used to simulate flux through *HK1* and *HK2* rich brain regions. A sensitivity analysis was conducted to determine how changes in K_m_ and basal intercellular glucose concentration effected the simulated change in glucose flux during hyperglycemia. Flux increased when K_m_ increased and when the basal intracellular glucose concentration decreased. Matrix axes are relative to the values for gray matter used by Barros et al. (K_m_ = 50 mM, glucose concentration = 1.2mM; (3)). **B)** Simulations from a single row, highlighted in white, in **A)** are shown over a range of blood glucose values. All flux values are percent change relative to euglycemic values. The light gray line is a simulation for gray matter and the dashed lines are the target blood glucose concentrations for euglycemia and hyperglycemia in this study. At lower basal brain glucose concentrations, the simulated change in flux through white matter was near the ∼60% value reported in Figure 2.

We then fixed the K_m_ of *HK2* to 5.6 times the K_m_ of *HK1* (50 mM; (3)), a value well within the range of 3 to 10 times reported in the literature (41). When the intracellular glucose concentration in white matter was a third of that in gray matter (1.2 mM; (3)), we found that acute hyperglycemia increased flux in white matter by ∼50% (**Figure 7B**). This is in general agreement with the 62% increase in white matter glucose consumption observed experimentally (**Figure 2C**). Conversely, flux through *HK1* in gray matter was insensitive to increases in blood glucose levels above concentrations of 5 mM. With higher euglycemic intracellular glucose concentrations, the change in white matter flux decreased, although an increase of ∼16% was still observed when the glucose concentration in white matter was equal to the gray matter concentration.

## Discussion

Several studies have examined the effects of acute hyperglycemia on regional cerebral glucose metabolism in humans (4–6,8,42). Our study is the first to examine changes in glucose metabolism together with regional changes in blood flow, blood volume, and oxygen consumption. Combining these measurements demonstrated that hyperglycemia elevates CMRglc in brain regions with low basal metabolic rates by increasing flux through non-oxidative pathways, or AG.

Although multiple studies have reported that acute hyperglycemia increases CMRglc in regions with low basal metabolic rates (4,5,43), there has been no attempt to explain this phenomenon. There is evidence that hyperglycemia increases excitability in hippocampal slices via potentiation of K_ATP_ channels (44). K_ATP_ channels are also necessary for the increase in interstitial fluid lactate, a putative marker of neuronal activity (45), in acutely hyperglycemic mice (12). Hyperglycemia increases the concentration of lactate in the blood (10) and multiple studies have shown that elevating serum lactate increases the cerebral blood flow response to sensory stimulation (38,46). Moreover, hyperglycemia is a well-known risk factor for seizures (47). Thus, it is possible to view increased CMRglc during hyperglycemia as an adaptive response to support elevated neuronal activity.

However, changes in neuronal activity normally are accompanied by proportional changes in CBF (36), which we did not observe during hyperglycemia in the task-free state. Nor did we observe augmented BOLD responses to task paradigms during hyperglycemia. Other investigators have also reported that hyperglycemia does not change the hemodynamic response to sensory stimulation in rats (48) or humans (49). More direct measurements (e.g., electrocorticography) are therefore needed to study the effects of hyperglycemia on neural activity in the human brain. In any case, our finding of elevated AG without a change in CBF or the BOLD response suggests that care should be taken when interpreting changes in cerebral lactate production as evidence of increased neuronal activity.

The strong regional correlation between basal CMRglc and change during hyperglycemia (**Figure 2D**; (4,43)) suggests that mechanisms that regulate CMRglc at rest are responsible for determining which regions are sensitive to hyperglycemia. Consistent with this hypothesis, we found that *HK2*, which has a K_m_ 3 to 10 times that of *HK1* (41), is highly expressed in white matter and other brain regions where hyperglycemia increased AG. Our simulations of glucose transport showed that, unlike *HK1*, *HK2* is not saturated at euglycemia and can accommodate increases in flux by upwards of 50% during hyperglycemia. The magnitude of the increase was inversely in proportion to the normal regional brain glucose concentration. Some investigators have found lower glucose concentrations in white matter than in gray matter (50), whereas others have reported similar (51) or higher values (2). Quantifying the precise contribution of *HK2* to increased glycolytic flux during hyperglycemia will require a more precise estimate of basal white matter glucose concentration. However, we note that, even if the concentration of glucose in white matter is higher than expected, there is evidence that glucose prevents the degradation of *HK2* (52), enabling increased flux even under saturated conditions (53).

Although some studies have reported little expression of *HK2* in the brain (41), two groups recently reported that *HK2* is expressed in microglia more than astrocytes, oligodendrocytes, or neurons in the mouse brain (54,55) (for a conflicting report see (56)). Interestingly, Hu et al found that *HK2* is upregulated in activated microglia in a number of disease models, implicating *HK2* in microglia function (54). Using two independent single-cell RNA sequencing datasets of the human brain (14,15), we were able to confirm that *HK2* expression is low in neurons and enriched in microglia and, to a lesser extent, astrocytes. Given the limited spatial resolution of PET, we cannot localize metabolic activity to a specific cell type. However, it is possible that our results are in part driven by microglia, as microglia density is higher in human white matter than gray matter (57), and tissue type influences microglia structure and function (58).

We can only speculate why the expression of *HK2* is elevated in white matter. It is rational to assume that hexokinases are expressed in relation to metabolic demand. Indeed, we found that the spatial distributions of the *HK1*/*HK2* ratio and resting glucose consumption are correlated. Therefore, *HK1* may be favored in gray matter as the lower K_m_ allows for greater enzymatic activity at typical blood glucose concentrations, and especially under conditions of hypoglycemia. Under conditions of euglycemia, the higher K_m_ of *HK2* makes it better suited for regions with lower metabolic rates. However, a higher K_m_ would be disadvantageous during hypoglycemia, which may explain, in part, the vulnerability of white matter to acute hypoglycemia (59).

Another possibility is that *HK2* is expressed in regions such as deep white matter that are susceptible to hypoxemia owing to their limited blood supply. This vulnerability manifests as deep white matter lesions that develop in a stereotypical peri-ventricular distribution in the setting of cerebrovascular small vessel disease (60) and sickle cell disease (61). Hypoxia increases the expression of *HK2* (62), a response mediated by hypoxia inducible factor (HIF) (63). During hypoxia, *HK2* binds to mitochondria, inhibiting apoptosis (62) induced by the creation of reactive oxygen species (64). Upregulation of *HK2* also enables glioblastoma cells to remain viable under hypoxic conditions at the tumor core (65). Thus, *HK2* exhibits multiple features that are protective against hypoxia.

Our finding of increased CMRglc without an increase in CMRO_2_ raises the question of determining which non-oxidative pathways increase flux during hyperglycemia. Lactate production is a strong possibility, as acute hyperglycemia elevates lactate levels in the interstitial fluid of the rodent hippocampus (12). However, studies in humans have not reported a significant increase in either cerebral lactate concentration (66) or efflux (67) during hyperglycemia.

Although glycogen production is another potential pathway, measuring changes in glycogen in the brain is challenging (68); and studies in rodents have reported both increased (69) and decreased (70) brain glycogen during acute hyperglycemia. Moreover, high intracellular glucose levels prompt the translocation of *HK2* from the cytoplasm, where it contributes to glycogen production (71), to the mitochondrial membrane, where it drives glycolysis (72). Acute hyperglycemia has been shown to increase activity through the pentose phosphate pathway (PPP) in cultured astrocytes (73). Activation of the PPP produces NADPH, conferring antioxidant benefits, and provides protection from amyloid-beta toxicity (74). Interestingly, the PPP is one of the mechanisms by which *HK2* reduces oxidative stress (75).

An intriguing possibly is that pathways upregulated in diabetes are responsible for increasing AG during acute hyperglycemia. Activation of the polyol pathway, which converts glucose to sorbitol and fructose, contributes to many of the clinical complications of diabetes, including cardiovascular disease, retinopathy, neuropathy, and cataracts (76). Although the polyol pathway plays a very minor role in the brain at euglycemia (77), both acute and chronic hyperglycemia increase polyol pathway activity in the brain (78,79). However, as the polyol pathway does use hexokinase to phosphorylate glucose, polyol activity cannot explain the correlation we found between *HK2* expression and CMRglc. Other prominent pathways that contribute to pathology in diabetes and could potentially increase AG include hexosamine synthesis (80) and the production of advanced glycated end products (81).

It has been reported that brain glucose transport is down-regulated in T2DM but these findings are controversial (for a review see (82)). It remains possible that chronically elevated glucose consumption contributes to white matter pathology in T2DM. Some studies have found that T2DM predisposes to white matter hyperintensities (83), white matter atrophy (84), and alterations in white matter microstructure (85). This perspective is adjacent to recent findings that upregulated glycolysis induced by increased expression of *HK2* is one of the mechanisms responsible for peripheral neuropathy in T2DM (86). Determining if hyperglycemia also increases AG in individuals with T2DM would be of considerable interest.

A few limitations should be considered in the interpretation of our results. First, although we did show that CMRglc quantitatively increases during hyperglycemia, we did not obtain quantitative measurement of oxygen consumption. Therefore, we were able only to show that oxygen consumption does not change relative to a reference region. However, because previous studies found no increase in global CMRO_2_ during acute hyperglycemia (67,87), we believe that the observed decrease in rOGI is most consistent with elevated AG. We do acknowledge, however, that relative PET techniques can be misleading (88). Thus, we defined a reference region by choosing brain regions with the smallest absolute change in CMRglc to minimize any potential confounds (**Figure S2**).

All of our measurements of CMRglc relied on [^18^F]FDG, which requires a correction factor, the lumped constant, to account for the differences in kinetics between FDG and glucose (89). Extrapolating from measurements in rats, we would predict that the lumped constant decreased by about 6% in our hyperglycemia data (90). As lower values of the lumped constant result in larger measured values of CMRglc, we potentially underestimated the increase in non-oxidative glucose consumption during hyperglycemia. To address this possibility, we followed the strategy of van Golen et al. (91), and adjusted the lumped constant for elevated blood glucose using the previously published rat data. Although this adjustment eliminated most of the significant decreases in CMRglc, it did not substantially alter the number of voxels with significantly elevated CMRglc (**Figure S2**). This is consistent with a previous study which reported elevated CMRglc during hyperglycemia using [1-^11^C]glucose (4). Given the minor impact of adjusting the lumped constant, we chose to use a fixed lumped constant in all primary analyses. However, the small decreases in CMRglc in **Figure 2**, as well as the rOGI increases in **Figure 4**, should be interpreted with caution given these findings.

Our hyperglycemic glucose clamp technique may have impacted our results. Specifically, we blocked the endogenous secretion of insulin and glucagon using octreotide and maintained glucagon and insulin at normal values throughout the experiment. Although this enables isolating the effect of hyperglycemia on brain metabolism, it is a somewhat non-physiological condition, as hyperglycemia normally sharply increases the concentration of insulin in the blood. However, as a defining feature of T2DM is insulin resistance, we believe that studying acute hyperglycemia without hyperinsulinemia increases our studies relevance to T2DM. It is also encouraging that our results are broadly consistent with prior studies that did not control insulin levels (4,6,42). Determining the effects of hyperglycemia with and without elevated insulin would be of considerable interest.

## Conclusion

Our work provides a novel addition to studies examining the effects of acute hyperglycemia on brain metabolism. We report that, in healthy adult humans, acute hyperglycemia increases CMRglc in white matter and other regions with low basal metabolic rates without altering regional blood flow, blood volume, or oxygen consumption. This result indicates that acute hyperglycemia increases AG in the absence of any evidence for increased neuronal activity. Among the most pressing outstanding issues are identifying the metabolic pathways responsible for elevated non-oxidative glucose use and determining if AG remains high in individuals with chronic hyperglycemia. Addressing these questions will further our understanding of brain metabolism during hyperglycemia and may clarify the relations between non-oxidative glucose consumption and T2DM.

## Acknowledgements

We thank all the participants that took part in this study. Pamela Stone, Kathleen Wharton, Angela Shackleford, Gleen Foster, and Linda Becker provided invaluable assistance in data collection. Drs. Beau Ances and Jiongjiong Wang shared the pCASL sequence. Dr. Pawel Markiewicz provided advice and access to NiftyPET in advance of its open-source release. Dr. Felipe Barros shared code for modeling glucose transport.

## Author contribution statement

TH, AMA, and MER designed research studies, JJL and AMA acquired data, JJL and TB analyzed data, and TB, AZS, MSG, and MER wrote the manuscript.

## Supplementary material

Supplementary material for this paper is available at: http://jcbfm.sagepub.com/content/by/supplemental-data

## Data availability

The data and code necessary to reproduce all of the figures and tables can be found at: https://doi.org/10.5281/zenodo.13647354. This repository includes individual-level regional averages for each imaging modality. Individual-level images will be made available upon reasonable request by a qualified researcher following a data usage agreement.

