## Supplemental Material for "Hyperglycemia selectively increases cerebral non-oxidative glucose consumption without affecting blood flow"

### Supplemental Information

| | $[^{18}F]$-FDG SUVR | | $[^{15}O]$-$H_2O$ SUVR | | $[^{15}O]$-$O_2$ SUVR | | rOEF SUVR | | $[^{15}O]$-CO SUVR | | rOGI SUVR | | Quant. CMRglc | | Quant CBF | | Task fMRI | |
|---|---|---|---|---|---|---|---|---|---|---|---|---|---|---|---|---|---|---|
| ID | Eugly. | Hyper. | Eugly. | Hyper. | Eugly. | Hyper. | Eugly. | Hyper. | Eugly. | Hyper. | Eugly. | Hyper. | Eugly. | Hyper. | Eugly. | Hyper. | Eugly. | Hyper. |
| sub-01 | ✓ | | ✓ | | ✓ | | ✓ | | ✓ | | ✓ | | | | | | | |
| sub-02 | ✓ | | ✓ | | ✓ | | ✓ | | ✓ | | ✓ | | | | | | | |
| sub-03 | ✓ | | ✓ | | ✓ | | ✓ | | ✓ | | ✓ | | | | | | | |
| sub-04 | ✓ | ✓ | ✓ | ✓ | ✓ | ✓ | ✓ | ✓ | ✓ | ✓ | ✓ | ✓ | | ✓ | ✓ | ✓ | ✓ | ✓ |
| sub-05 | ✓ | | ✓ | | ✓ | | ✓ | | ✓ | | ✓ | | | | ✓ | | | |
| sub-06 | ✓ | ✓ | ✓ | ✓ | ✓ | ✓ | ✓ | ✓ | ✓ | ✓ | ✓ | ✓ | | | ✓ | ✓ | ✓ | ✓ |
| sub-07 | ✓ | ✓ | ✓ | ✓ | ✓ | ✓ | ✓ | ✓ | ✓ | ✓ | ✓ | ✓ | | | ✓ | ✓ | ✓ | ✓ |
| sub-08 | ✓ | ✓ | ✓ | ✓ | ✓ | ✓ | ✓ | ✓ | ✓ | ✓ | ✓ | ✓ | | | ✓ | ✓ | ✓ | ✓ |
| sub-09 | | ✓ | ✓ | ✓ | ✓ | ✓ | ✓ | ✓ | ✓ | ✓ | | ✓ | | | ✓ | ✓ | ✓ | ✓ |
| sub-10 | ✓ | | ✓ | | ✓ | | ✓ | | ✓ | | ✓ | | | | ✓ | | | |
| sub-11 | ✓ | ✓ | ✓ | ✓ | ✓ | ✓ | ✓ | ✓ | ✓ | ✓ | ✓ | ✓ | | | ✓ | ✓ | ✓ | ✓ |
| sub-12 | ✓ | ✓ | ✓ | ✓ | ✓ | ✓ | ✓ | ✓ | ✓ | ✓ | ✓ | ✓ | | | ✓ | ✓ | ✓ | ✓ |
| sub-13 | ✓ | ✓ | ✓ | ✓ | ✓ | ✓ | ✓ | ✓ | ✓ | ✓ | ✓ | ✓ | ✓ | ✓ | ✓ | ✓ | ✓ | ✓ |
| sub-14 | ✓ | ✓ | ✓ | ✓ | ✓ | ✓ | ✓ | ✓ | ✓ | ✓ | ✓ | ✓ | ✓ | ✓ | ✓ | ✓ | ✓ | ✓ |
| sub-15 | ✓ | ✓ | ✓ | ✓ | ✓ | ✓ | ✓ | ✓ | ✓ | ✓ | ✓ | ✓ | ✓ | ✓ | ✓ | ✓ | ✓ | ✓ |
| sub-16 | ✓ | ✓ | ✓ | ✓ | ✓ | ✓ | ✓ | ✓ | ✓ | ✓ | ✓ | ✓ | ✓ | | ✓ | ✓ | ✓ | |
| sub-17 | ✓ | | ✓ | | ✓ | | ✓ | | ✓ | | ✓ | | | | ✓ | | ✓ | |
| sub-18 | ✓ | ✓ | ✓ | ✓ | ✓ | ✓ | ✓ | ✓ | ✓ | ✓ | ✓ | ✓ | ✓ | ✓ | ✓ | ✓ | ✓ | ✓ |
| sub-19 | ✓ | ✓ | ✓ | ✓ | ✓ | ✓ | ✓ | ✓ | ✓ | ✓ | ✓ | ✓ | ✓ | ✓ | ✓ | ✓ | ✓ | ✓ |
| sub-20 | | | | | | | | | | | | | | | ✓ | | ✓ | |
| sub-21 | ✓ | ✓ | ✓ | ✓ | ✓ | ✓ | ✓ | ✓ | ✓ | ✓ | ✓ | ✓ | ✓ | ✓ | ✓ | ✓ | ✓ | ✓ |
| sub-22 | ✓ | ✓ | ✓ | ✓ | ✓ | ✓ | ✓ | ✓ | ✓ | ✓ | ✓ | ✓ | ✓ | ✓ | ✓ | ✓ | ✓ | ✓ |
| sub-23 | ✓ | ✓ | ✓ | ✓ | ✓ | ✓ | ✓ | ✓ | ✓ | ✓ | ✓ | ✓ | ✓ | ✓ | ✓ | ✓ | ✓ | ✓ |
| sub-24 | | ✓ | | ✓ | | ✓ | | ✓ | | ✓ | | ✓ | | ✓ | | ✓ | | ✓ |
| sub-25 | ✓ | ✓ | ✓ | ✓ | ✓ | ✓ | ✓ | ✓ | ✓ | ✓ | ✓ | ✓ | ✓ | ✓ | ✓ | ✓ | ✓ | ✓ |
| sub-26 | ✓ | ✓ | ✓ | ✓ | ✓ | ✓ | ✓ | ✓ | ✓ | ✓ | ✓ | ✓ | ✓ | ✓ | ✓ | ✓ | ✓ | ✓ |
| sub-27 | ✓ | ✓ | ✓ | ✓ | ✓ | ✓ | ✓ | ✓ | ✓ | ✓ | ✓ | ✓ | ✓ | ✓ | ✓ | ✓ | ✓ | ✓ |
| sub-28 | | ✓ | | ✓ | | ✓ | | ✓ | | ✓ | | ✓ | | ✓ | | ✓ | | ✓ |
| sub-29 | ✓ | | ✓ | | ✓ | | ✓ | | ✓ | | ✓ | | ✓ | | ✓ | | ✓ | |
| Totals | 25 | 21 | 26 | 21 | 26 | 21 | 26 | 21 | 26 | 21 | 25 | 21 | 13 | 14 | 24 | 21 | 22 | 20 |

**Supplemental Table 1:** Breakdown of participants contributing to each imaging datatype.

| Time | Glucose (mg·dL$^{-1}$·min$^{-1}$) | | Insulin (pmol·L$^{-1}$·min$^{-1}$) | |
|---|---|---|---|---|
| | **Eugly.** | **Hygly.** | **Eugly.** | **Hygly.** |
| **Before breakpoint** | 0.38 ± 0.1 | 3.9 ± 0.11 | 0.72 ± 0.24 | 0.59 ± 0.28 |
| **After breakpoint** | -0.06 ± 0.02 | -0.14 ± 0.02 | -0.13 ± 0.063 | 0.35 ± 0.07 |

**Supplemental Table 2:** The concentrations of plasma glucose and insulin during our study were not entirely steady after the desired plasma glucose level was reached (**Figure 1**). To quantify these observations, we performed a piecewise linear regression where the slope of the regression line was allowed to differ after 55 minutes into the glucose clamp (see **Methods**). Slope estimates from the piecewise regression are shown above**.** Values are means and symmetric 95% confidence intervals. All slopes are significantly different from zero at the 0.05 level without correction for multiple comparisons.

For the first 55 minutes, plasma glucose and insulin increased with time in both conditions ($p < 0.0001$). However, after 55 minutes plasma glucose decreased with time during both clamps ($p < 0.0001$), and plasma insulin decreased in the euglycemic condition ($p < 0.0001$). Plasma insulin continued to increase slightly with time after 55 minutes during the hyperglycemic clamp ($p < 0.0001$).

| Parameter | Eugly. | Hygly. | Hygly. – Eugly. | p-value |
|---|---|---|---|---|
| **$K_1$ ($mL \cdot hg^{-1} \cdot min^{-1}$)** | 12.67 ± 1.01 | 6.61 ± 1.00 | -6.06 ± 0.92 | **$5.18 \cdot 10^{-8}$** |
| **$k_2$ ($min^{-1}$)** | 0.31 ± 0.042 | 0.22 ± 0.041 | -0.091 ± 0.47 | **$3.24 \cdot 10^{-3}$** |
| **$k_3$ ($min^{-1}$)** | 0.17 ± 0.021 | 0.072 ± 0.021 | -0.096 ± 0.029 | **$4.8 \cdot 10^{-5}$** |
| **$k_4$ ($min^{-1}$)** | 0.014 ± 0.003 | 0.026 ± 0.003 | 0.012 ± 0.004 | **$6.17 \cdot 10^{-5}$** |
| **$V_b$ ($mL \cdot hg^{-1}$)** | 4.86 ± 0.63 | 4.66 ± 0.62 | -0.20 ± 0.85 | $6.51 \cdot 10^{-1}$ |
| **CMRglc ($\mu Mol \cdot hg^{-1} \cdot min^{-1}$)** | 30.70 ± 1.90 | 33.54 ± 1.88 | 2.84 ± 2.41 | **$4.15 \cdot 10^{-2}$** |

**Supplemental Table 3:** Whole-brain parameter estimates from the reversible 2-compartment FDG model. For each parameter, a linear mixed model was used to estimate the population mean and 95% CI during both the euglycemic (n=13) and hyperglycemic (n=14) clamp. $K_1$, $k_2$, and $k_3$ significantly decreased during hyperglycemia, whereas $k_4$ and CMRglc (cerebral metabolic rate of glucose consumption) significantly increased ($p < 0.05$; bold text).

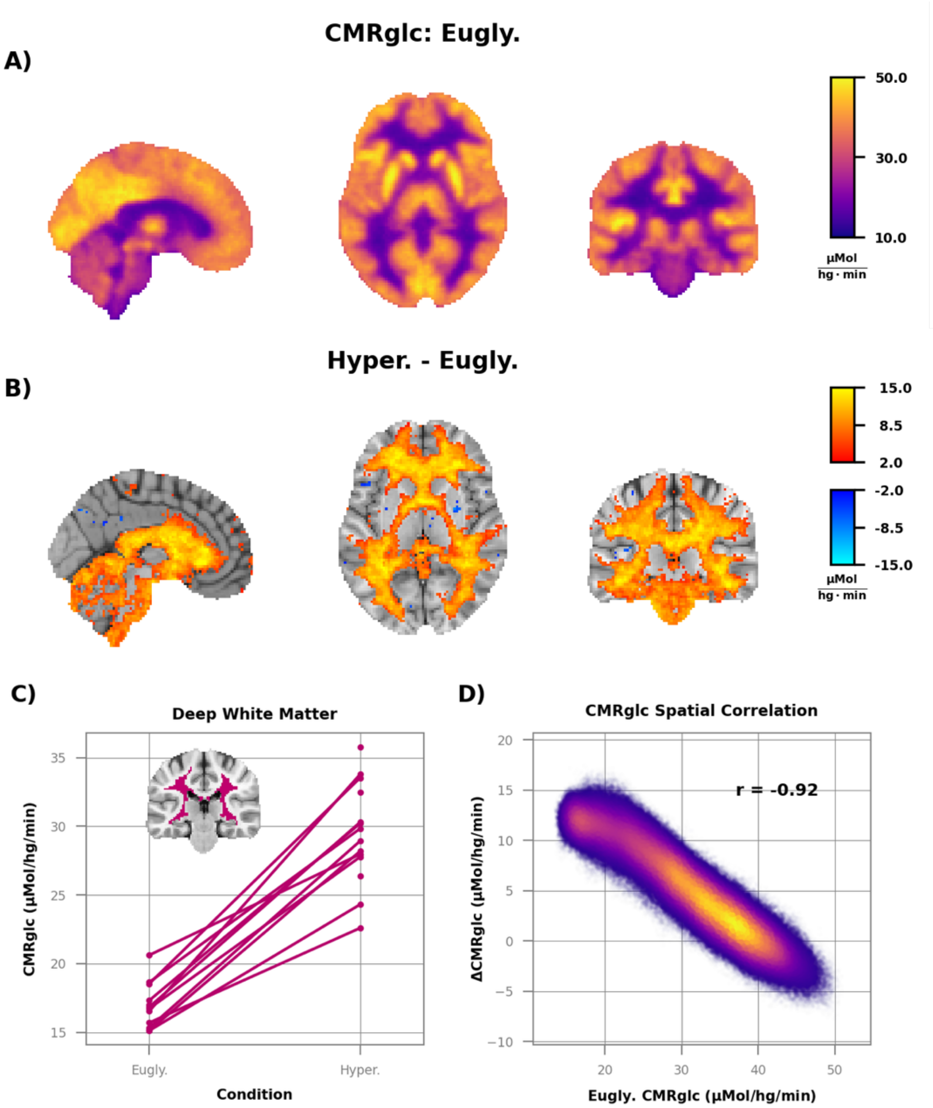


**Supplemental Figure 1:** Quantification of CMRglc with PET requires a corrected factor to account for differences in transport between [$^{18}$F]FDG and glucose. This correction factor is historically called the lumped constant. There is evidence that the lumped constant decreases slightly during hyperglycemia (1). To determine what impact changes in the lumped constant could have on our results, we computed the values shown in **Figure 2** after adjusting for potential differences in the lumped attributable to glycemic level (see **Methods**). All figures follow the same conventions as **Figure 2**. The focal increases in CMRglc were reported in Figure 2 were not affected by the lumped constant adjustment. However, the decreases in CMRglc that were found throughout gray matter in **Figure 2** were greatly diminished after this correction.

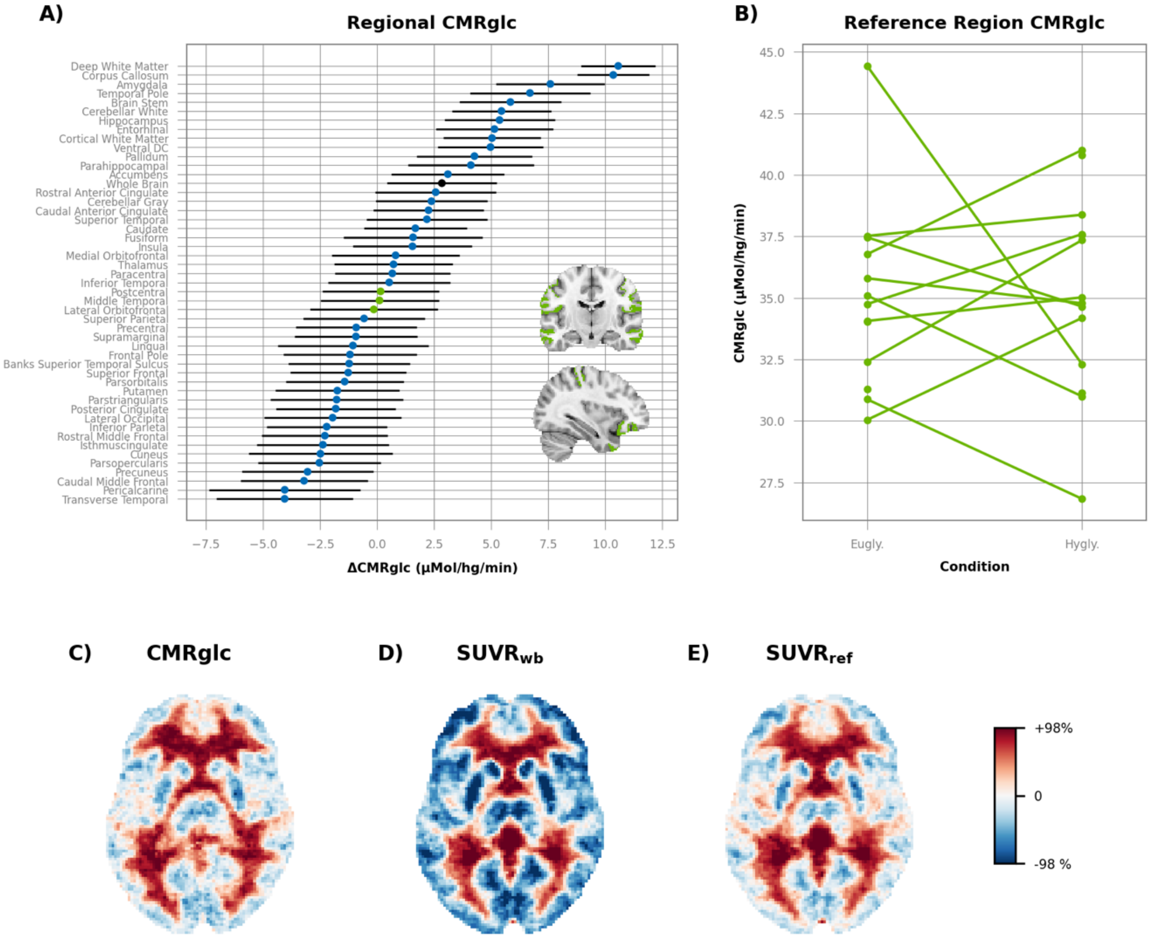


**Supplemental Figure 2:** Identification of a reference region for relative PET SUVR measurements**. A)** Group-average CMRglc for 48 non-overlapping brain regions defined by FreeSurfer. Error bars are 95% confidence intervals. The black dot is the whole brain average, and the green dots are the three regions with the smallest absolute change in CMRglc during hyperglycemia (middle temporal, lateral orbitofrontal, and postcentral gyrus). For PET SUVR measurements these three regions were combined into a single reference region. **B)** CMRglc within the combined reference region ROI. There was no significant difference between conditions ($0.06 \pm 2.62$ $\mu Mol \cdot hg^{-1} \cdot min^{-1}$; $p = 0.96$). Group average images of the difference in glucose consumption between hyperglycemia and euglycemia **using C)**

quantitative CMRglc (n = 27 scans), **D)** whole-brain SUVR (n=46 scans), and **E)** SUVR using the combined reference region (n=46 scan). The maximum colormap value was matched to the 98% percentile value for each image to account for the difference in scale between the CMRglc and SUVR images. Because hyperglycemia increased the whole-brain CMRglc slightly (see **Results**), using a whole-brain reference region produced larger decreases in relative gray matter glucose consumption (**D**) than was seen with quantitative CMRglc (**C**). Using the empirically defined combined reference region (**E**) produced results much more consistent with quantitative CMRglc. Note that the SUVR data in **D**) and **E**) was not spatially smoothed in order to match the CMRglc data in **C**).

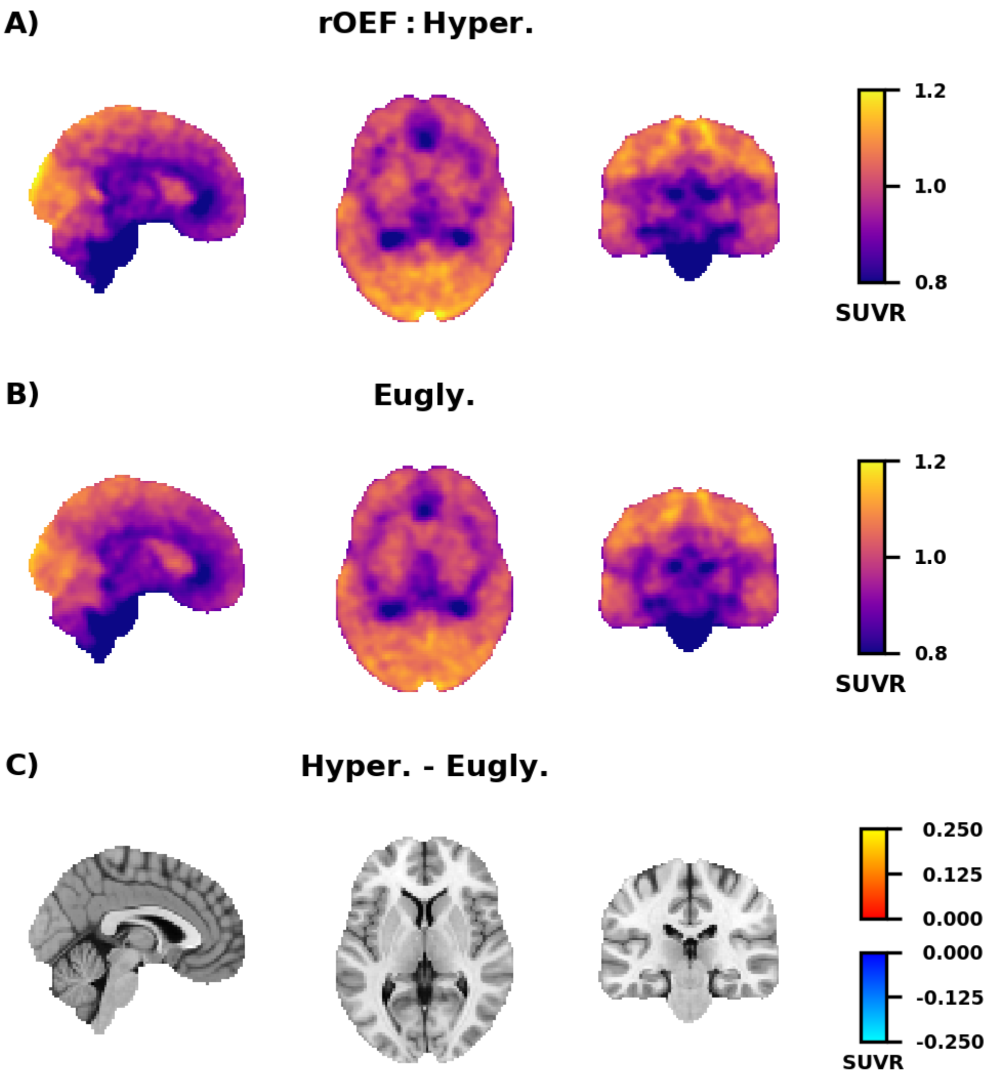


**Supplemental Figure 3**: **A)** Group average (n=26) image of the relative oxygen extraction (rOEF) measured [$^{15}$O]$O_2$ and [$^{15}$O]$H_2O$ SUVR. Values are normalized to an empirically derived reference region (see **Supplemental Figure 2**). **B)** Group average (n=21) image rOEF during the hyperglycemic clamp. **C)** Group average difference in rOEF between the hyperglycemic and euglycemic clamp. Only voxels that are significantly different from zero after correction for multiple comparisons (False Discovery Rate 0.05) are shown in color. Voxels where rOEF decreased during hyperglycemia are in blue, whereas increases are shown in orange/yellow.

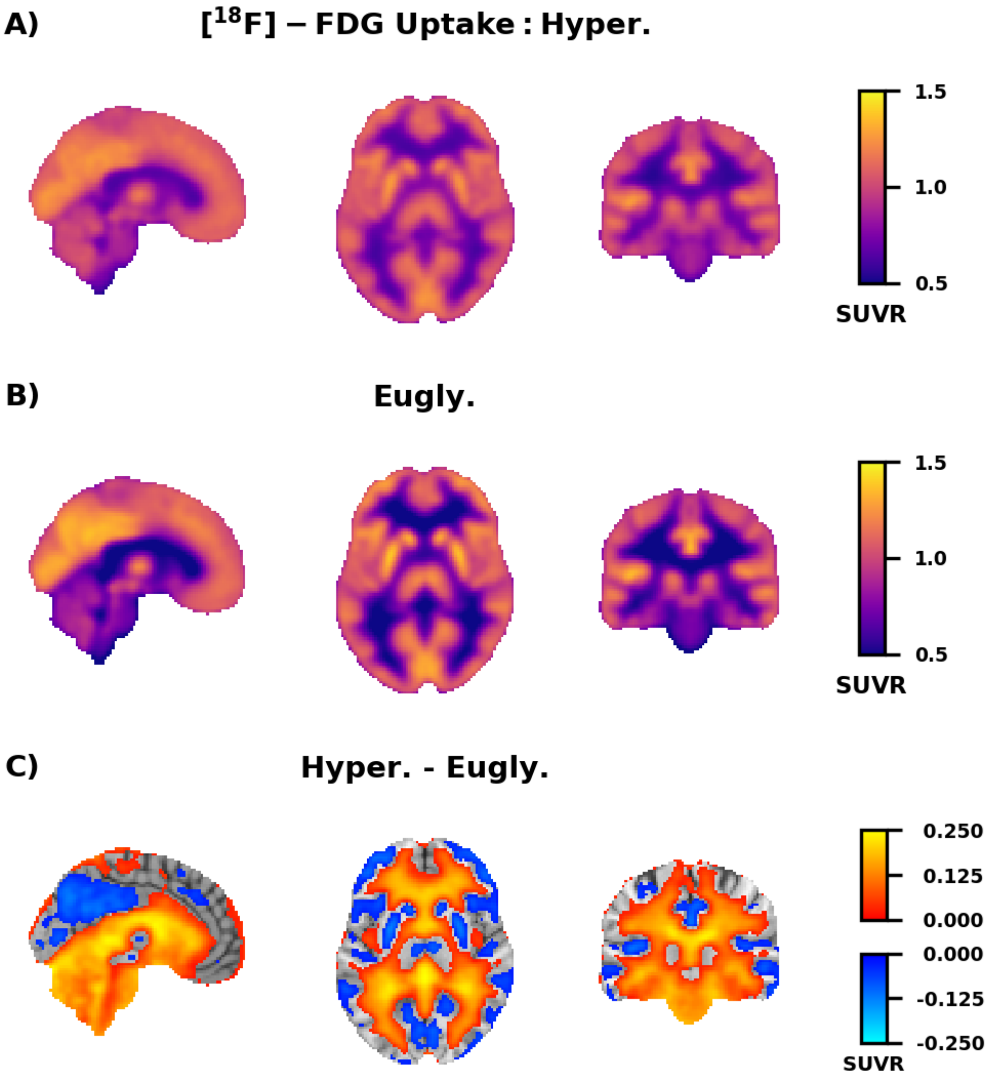


**Supplemental Figure 4:** Hyperglycemia induced changes in relative glucose consumption measured with normalize [$^{18}$F]FDG SUVR. All conventions as **Supplemental Figure 3**. As in **Figure 2**, glucose consumption was significantly ($p$ < 0.05) greater in the white matter during **B)** hyperglycemia (n=21) than **A)** euglycemia (n=25).

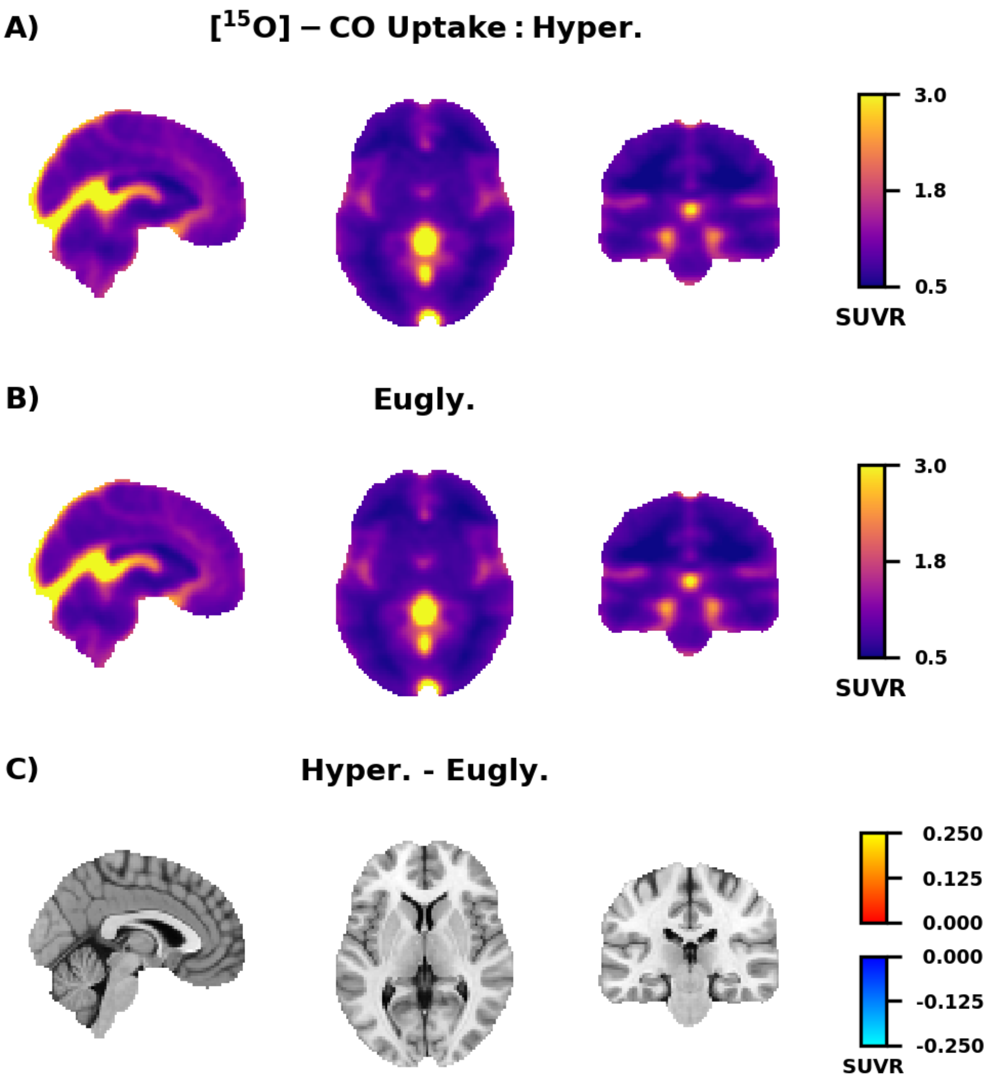


**Supplemental Figure 5.** Relative cerebral blood volume measured with normalized $[^{15}O]CO$ SUVR. All conventions as in Figure S3. No significant differences ($p < 0.05$) were found between **A)** euglycemia (n=25) and **B)** hyperglycemia (n=21) after **C)** correction or multiple comparisons with the False Discovery Rate.

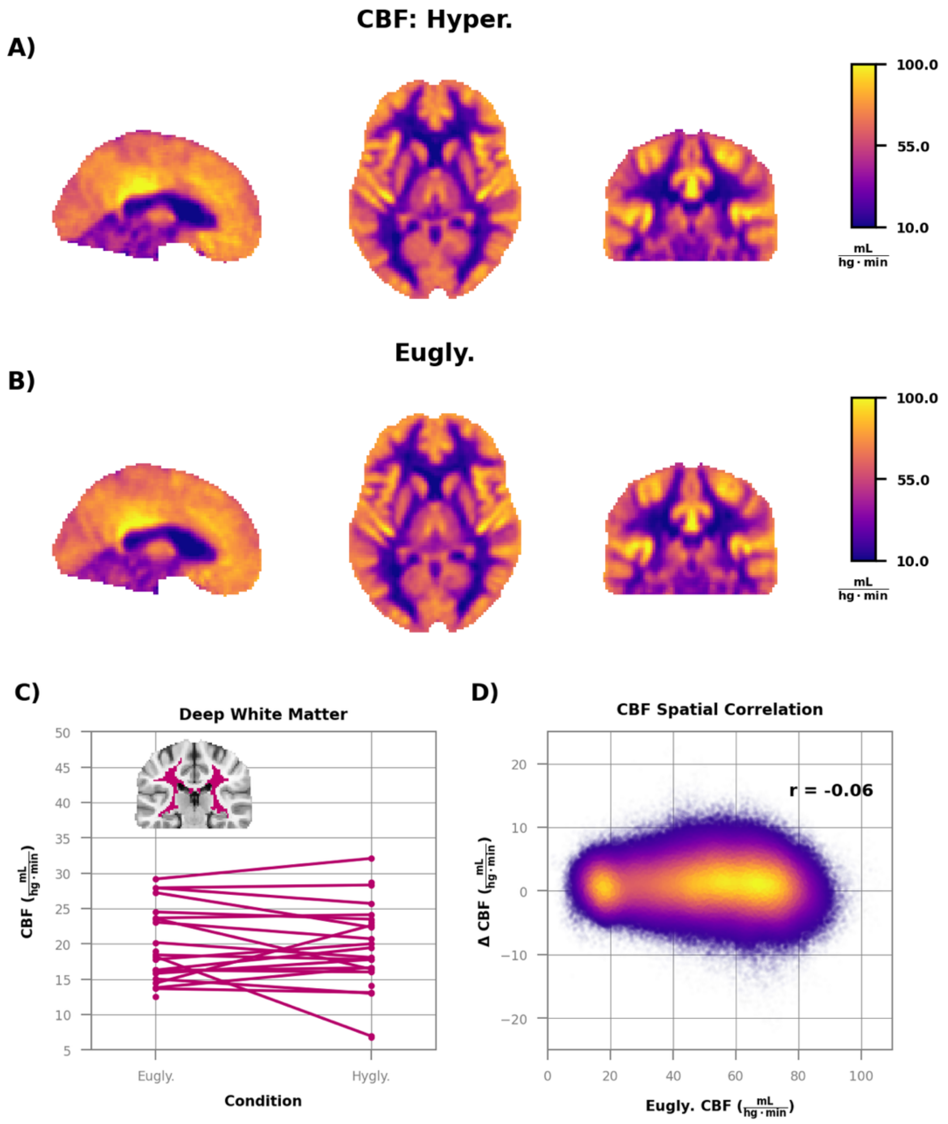


**Supplemental Figure 6:** No change in pCASL MRI derived quantitative CBF during hyperglycemia. The group average of images CBF during **A)** hyperglycemia (n=21) and **B)** euglycemia (n=27) are essentially identical. No statistically significant differences *($p$* $< 0.05$; False Discovery Rate corrected) were found between the two conditions. **C)** CBF within the deep white matter. The average difference between conditions ($0.51 \pm 1.81$ mL·hg$^{-1}$·min$^{-1}$) was not significant *($p$* $= 0.58$). **D)** Scatterplot of CBF during euglycemia vs. the difference between hyperglycemia and euglycemia. Each point is a voxel and color indicates density. There was no relationship between baseline CBF and hyperglycemia induced change.

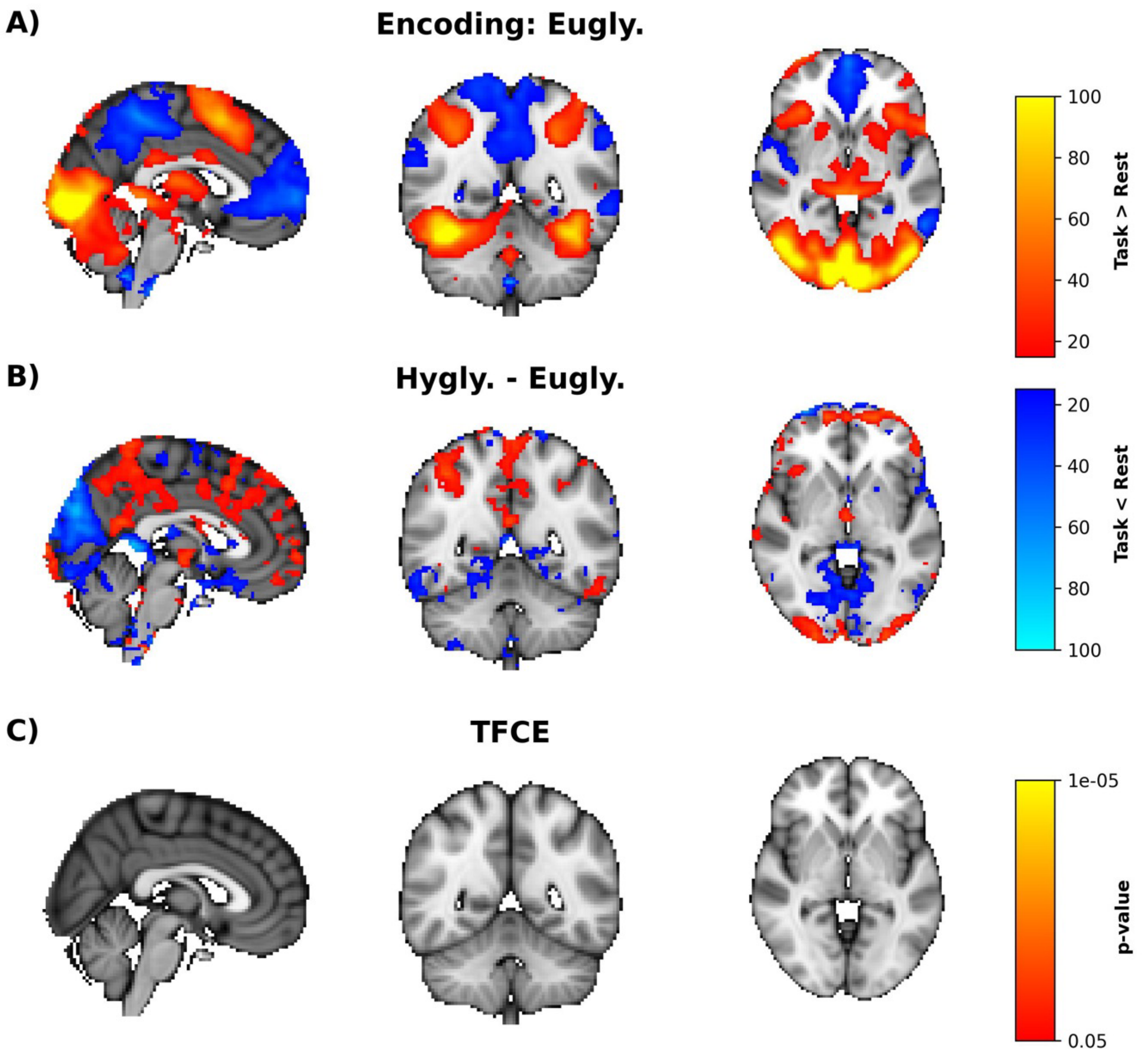


**Supplemental Figure 7:** Hyperglycemia does not affect activity evoked by a face-name encoding task. **A)** Group average image of brain regions activated (red/yellow) and deactivated (blue/cyan) by the task. **B**) Average difference in BOLD activity between hyperglycemia (n=20) and euglycemia (n=22). **C)** None of the differences in **B)** were significant after correction for multiple comparisons using threshold-free cluster enhancement.

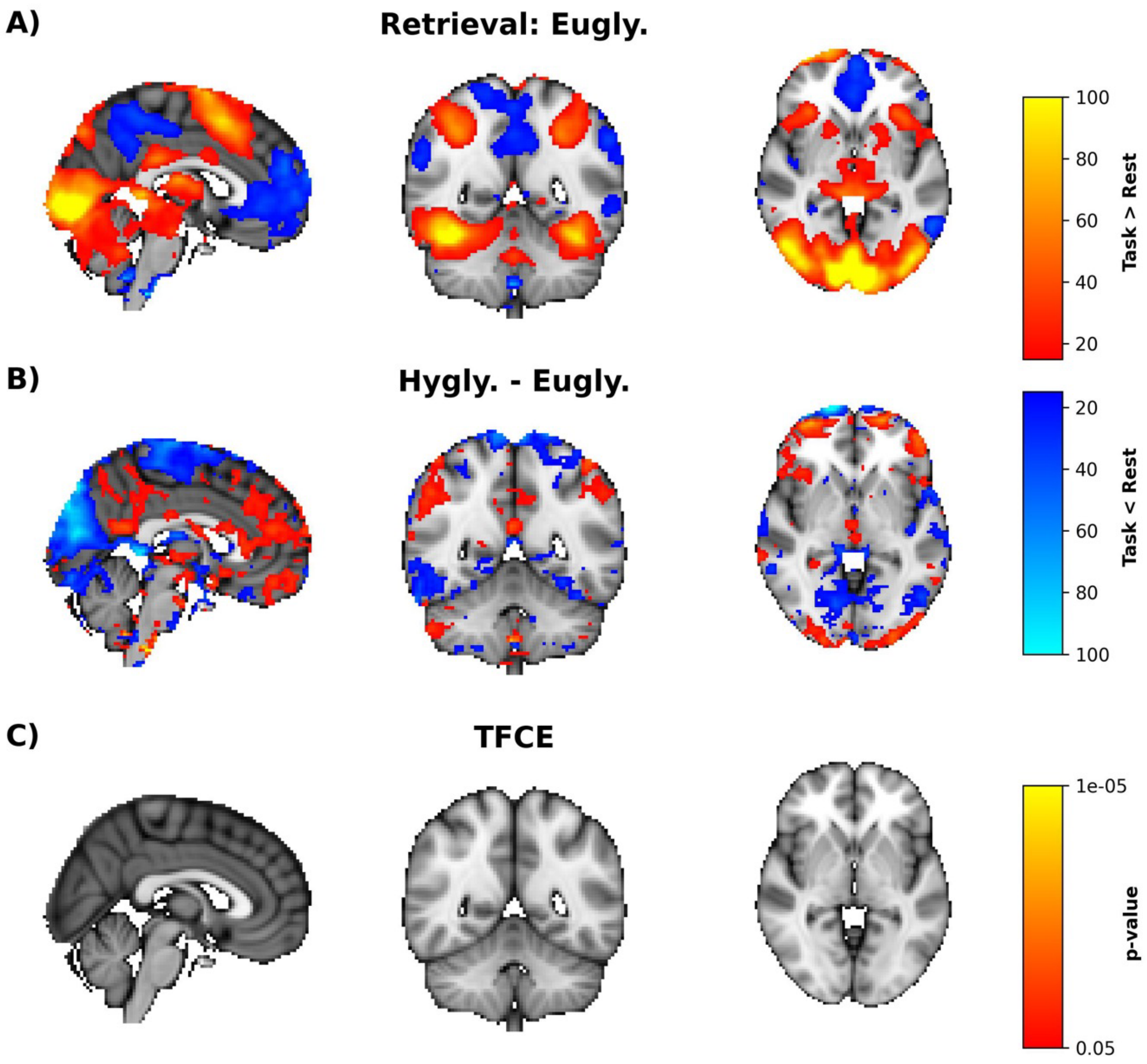


**Supplemental Figure 8:** Hyperglycemia does not affect activity evoked by a face-name retrieval task. **A)** Group average image of brain regions activated (red/yellow) and deactivated (blue/cyan) by the task. **B**) Average difference in BOLD activity between hyperglycemia (n=20) and euglycemia (n=21). **C)** None of the differences in **B)** were significant after correction for multiple comparisons using threshold-free cluster enhancement.

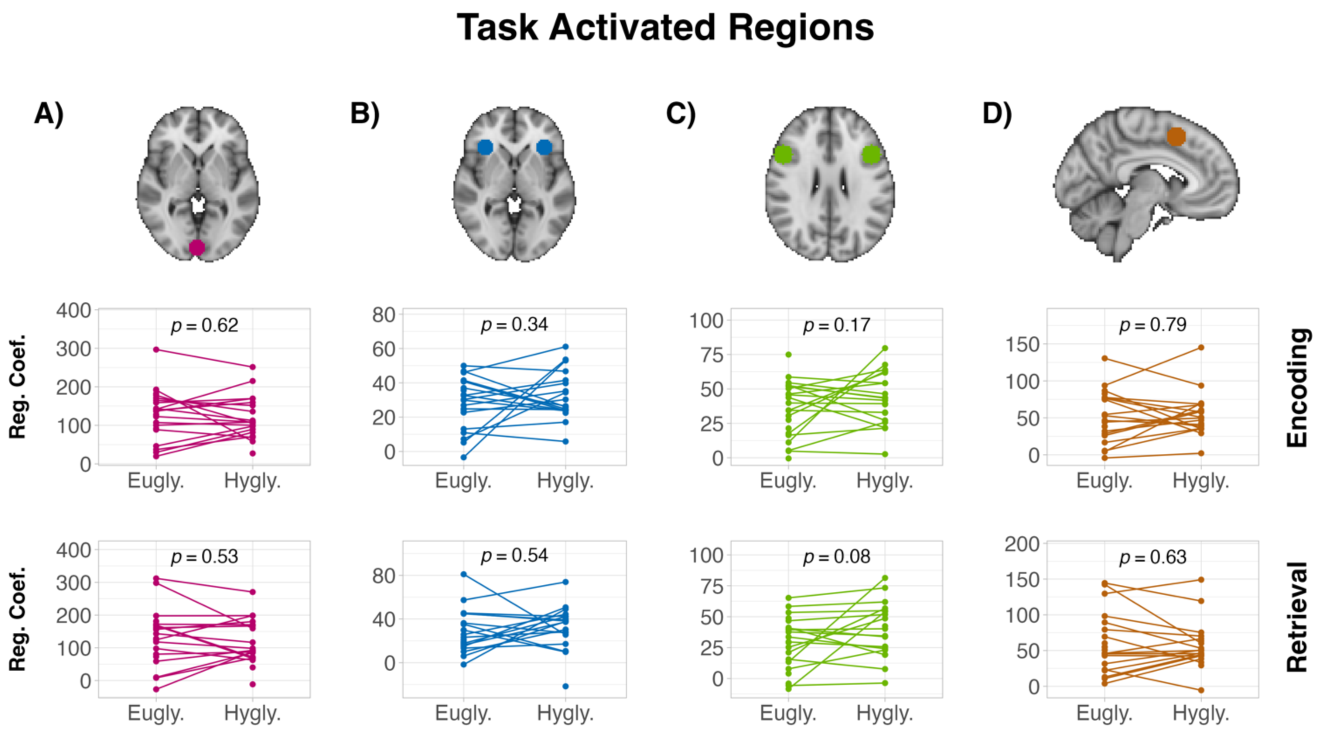


**Supplemental Figure 9:** Four ROIs were defined by placing 10 mm spheres on the task-activated regions in **Supplemental Figure 7**. Average responses within the ROIs are plotted for the encoding task (second row) and the retrieval task (third row). No significant differences were found in any of the ROIs in either condition. However, there was a trend-level increase in the inferior frontal gyrus during retrieval **C**).

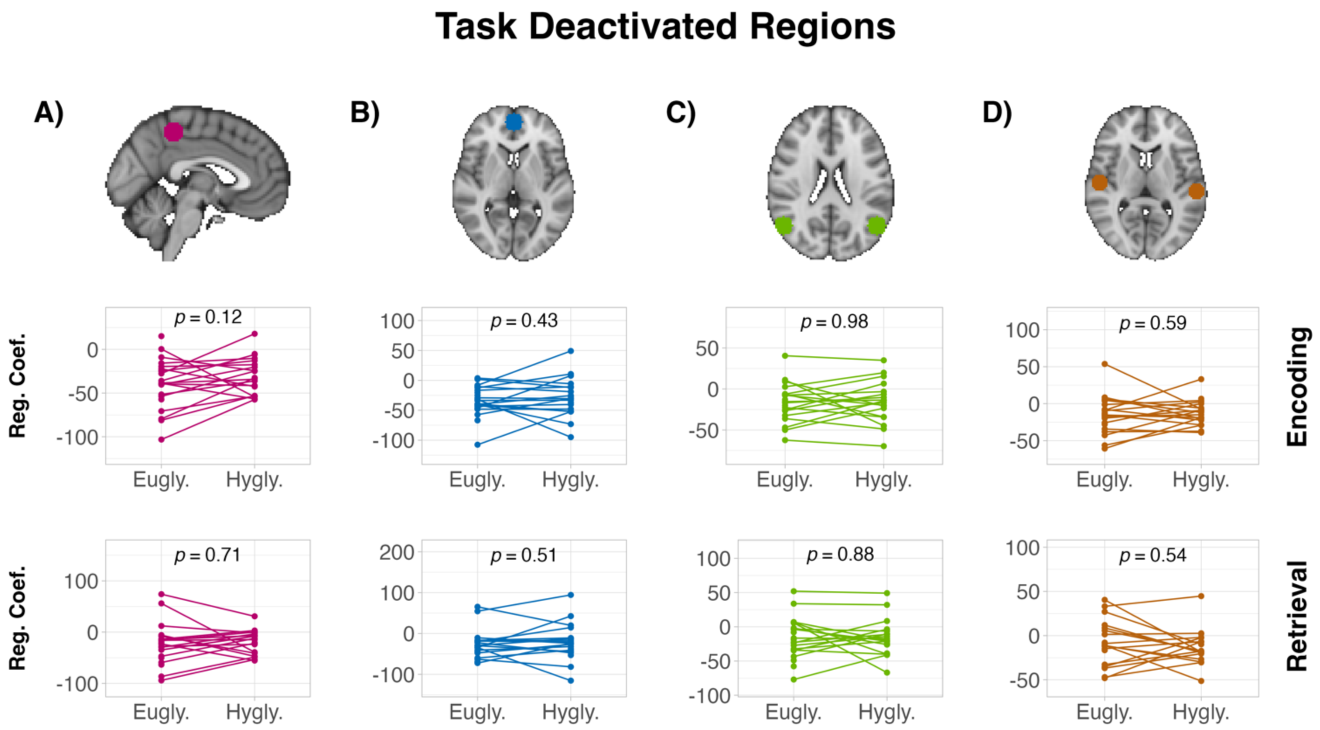


**Supplemental Figure 10:** Four ROIs were defined by placing 10 mm spheres on the task-deactivated regions in **Supplemental Figure 7**. Average responses within the ROIs are plotted for the encoding task (second row) and the retrieval task (third row). No significant differences were found in any of the ROIs in either condition. However, there was a trend-level increase (i.e., decrease in deactivation) in the precuneus during encoding **A**).

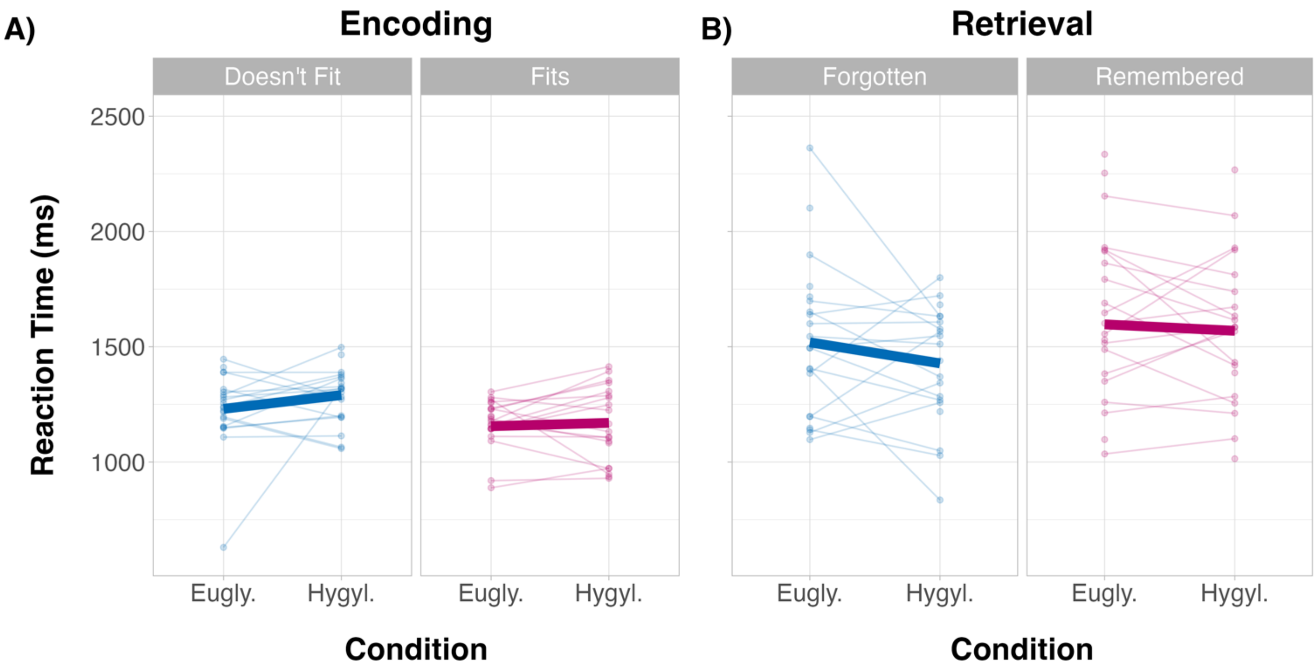


**Supplemental Figure 11:** No effect of hyperglycemia on reaction time. **A)** During the encoding condition participants were shown a face-name pair and asked whether the name fit the face. There was no difference in reaction time between euglycemia and hyperglycemia for either response ($p > 0.05$). **B)** In the retrieval condition participants were shown an image of a face and asked if they remembered the face. Again, hyperglycemia did not change reaction time ($p > 0.05$).

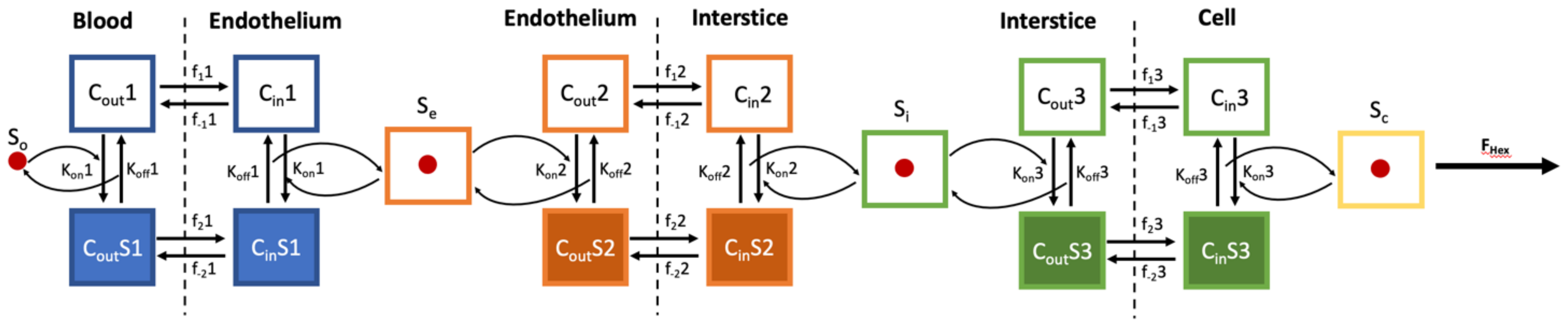


**Supplemental Figure 12:** Model of glucose transport introduced by Barros et al. (2). Filled boxes are carriers with bound glucose and empty boxes are unbound carriers. Dashed lines represent membrane boundaries. $C_{out}$ and $C_{in}$ are the unbound carrier concentrations on either side of the membrane, while $C_{out}S$ and $C_{in}S$ are the corresponding values for carriers bound with glucose. Rates constants for movement of the carrier across the membrane are denoted with an $f_x n$ where x=1 denotes a unbound carrier, x=2 a bound carrier, and n indicates the membrane (1, 2, or 3). $K_{off}n$ and $K_{on}n$ are the dissociation and association constants for each membrane. $S_o$, $S_e$, $S_i$, and $S_c$ are the concentration of glucose in the blood, endothelium, interstice, and intracellular comparents respectively. $F_{hex}$ is the irreversible breakdown of glucose. See **Methods** for an explanation of the simplifications that were performed to facilitate solving the system of differential equations.

**Supplemental Methods:**

Glucose clamps

Participants were admitted in the morning after a 10-hour overnight fast. Two intravenous catheters were inserted into antecubital veins (one for infusion of octreotide, glucagon, insulin, dextrose and potassium, and the other line for injection of radioactive isotopes. A radial artery was cannulated for arterial sampling of analytes (glucose and insulin) every 30 minutes during the clamps and for radioactivity sampling after the injection of the radiotracer. Blood pressure and heart rate was recorded every 30 minutes and cardiac function was monitored by electrocardiography throughout the study.

We infused the somatostatin analog octreotide (Teva Pharmaceuticals USA, Inc. North Wales, Pennsylvania) in a dose of 30 $ng·kg^{-1}·min^{-1}$ in all subjects to suppress endogenous insulin and glucagon secretion (3). The suppression of endogenous insulin was critically important to isolate the effect of hyperglycemia on brain metabolism. Basal insulin concentrations were ensured by infusing insulin at a rate of 0.1 $mU·kg^{-1}·min^{-1}$ (Eli Lilly, Indianapolis, Indiana). Basal glucagon was replaced using a glucagon dose of 1.0 $ng·kg^{-1}·min^{-1}$ (Fresenius Kabi Lake Zurich, Illinois). The dose rates for insulin and glucagon were determined by previous studies conducted by our group (4). Hormone infusates were prepared in saline containing 0.5 $g·dL^{-1}$ albumin. Potassium chloride was given at a rate of 5 $mmol·h^{-1}$ to prevent insulin-induced hypokalemia (5). 20% dextrose was infused intravenously at a variable rate based on bedside plasma glucose determinations (YSI Glucose Analyzer 2, Yellow Springs Instruments, Yellow Springs, OH) every five to ten minutes, to keep plasma glucoses levels clamped at euglycemia (90-100 $mg·dL^{-1}$ ) or hyperglycemia (250-300 $mg·dL^{-1}$).

The euglycemic clamp was performed first in 22 of the participants . Three participants had data from two separate euglycemic clamp visits. To account for the correlation between repeat visits, we used linear mixed models when performing statistical analyses . The second euglycemic visit was acquired as part of a separate study exploring the effect of hyperinsulinemia on brain metabolism.

pCASL MRI preprocessing

Images were then motion corrected using FSL's (6) MCFLIRT tool with the temporal mean image as the target (7). Following the recommendation of Wang et al., motion correction was performed separately for label and control frames (8). After motion correction, FSL's FLIRT was used to compute a rigid body transformation between the average label and average control images. A rigid body transformation was then computed between the realigned pCASL timeseries and each subject's average T1-weighted image using FSL's flirt. For subjects where field maps were acquired, the field map magnitude image was used as an intermediary between the pCASL and T1-weighted images. Nonlinear registration between T1-weighted image and MNI152 space was performed using FNIRT in FSL (6)._All transformations were then composed, and the pCASL time series was resampled to MNI152 2mm atlas space in a single step.

When computing the mean CBF for a single pCASL run, a weighting scheme was applied to down weight frames where subject motion produced large global shifts in image intensity (9). The mean difference in global signal between control and label images (-20.19 ± 77.49) between hyperglycemic (262.46 ± 60.14) and euglycemic (282.65 ± 55.39) runs was not significantly different from zero ($p$ = 0.62). The median CBF for each session was calculated over the whole brain using all available voxels.

Task MRI preprocessing

Runs whose average absolute or relative movement was greater than two standard deviations from the mean (absolute = 1.5 mm, relative = 0.36 mm) were excluded from all analyses. When field map images were available, geometric distortions in the phase encoding direction were removed using the FUGUE and PRELUDE tools (10). Rigid-body registration with a gray/white boundary-based cost function (11) was used to align the EPI images to each participant's T1-weighted anatomical image. FSL's FNIRT was used to compute a nonlinear transformation between each participant's anatomical image and MNI152 space.

Each task run was modeled as events with a 2-second duration convolved with a double gamma hemodynamic response function intermixed with jittered periods of rest. The temporal derivative of this regressor was also included in the model. Runs within a subject and task (e.g., both encoding runs from one day for one subject) were combined using a fixed-effects model in FILM (12). To mitigate the effect of motion, fsl_motion_outliers was used to identity high motion frames. A frame was flagged as an outlier if the root-mean-square (RMS) difference between it and first frame in the run was greater than third quartile plus 1.5 times the interquartile range. A separate regressor for each outlier frame was added to the fixed effect model. After removing outlier frames there was no difference in average frame-to-frame mean squared error between hyperglycemia and euglycemia during either encoding ($p$=0.12) or retrieval ($p$=0.07).

CT imaging

For attenuation correction, a Siemens Biograph 40 PET/CT was used to acquire a CT image of the head (120 keV, 25 effective mAs, voxel size = 0.59 x 0.59 x 3.0 mm, acquisition matrix = 512 x 512 x 74 mm voxels). From the CT image, a μ-map was created by converting the CT Hounsfield values into attenuation coefficients (13). Subject-specific μ-maps were combined with PET/MR hardware μ-maps provided by the manufacturer (Siemens).

PET preprocessing

Dynamic image frames were generated using axial compression, four iterations of ordered sets with expectation maximization, Gaussian filtering to 4.3 mm FWHM, randoms corrections by delayed timing, and a novel voxel-based single scatter model implementing 3D scatter sinograms (33). To ensure the highest possible data quality, our PET reconstruction strategy consisted of two stages. In the first stage, the listmode data were scatter corrected and reconstructed without attenuation or scatter correction using frame durations: 12, 13, 14, 15, 17, 18, 20, 23, 26, 30, 35, 43, 55, 75 and 114 seconds. [$^{18}$F]FDG images were reconstructed using frame durations of: 30, 35, 39, 43, 47, 51, 55, 59, 64, 68, 72, 76, 81 , 85, 89, 93, 98, 102, 106, 111, 115, 120, 124, 129, 133, 138, 142, 147, 151, 156, 161, 165, 170, and 175 seconds. Attenuation correction was not done at this stage because motion between frames precluded the use of a single μ-map for all frames.

In the second stage, we used a modified version of a previously published strategy to correct for between-frame motion (34). Briefly, for each scan each frame was registered to every other frame. From this set of pairwise registrations, a linear system of transformation cycles was created from which it was possible to compute the least squares estimate of rigid body transformations between any two frames. These transformations were used to align the previously computed μ-map with the time-resolved PET reconstructions. The aligned μ-maps were used in the second stage of reconstruction to create time-sliced PET images with attenuation, decay, and scatter and additional motion correction. The use of attenuation and scatter correction in the second stage allowed us to use shorter time bins for PET reconstruction. The first frame for the final reconstructed $^{15}$O image began up three seconds prior to tracer inflow. Subsequent frames had progressively increasing durations which were inversely proportional to rate of radionucleotide decay (λ): $\Delta \sim 2^{\frac{t}{\lambda}}$. Because of variable inflow times, the number and duration of frames was variable across scans. The frame durations for [$^{18}$F]FDG increased by shorter proportional increments, beginning with a 10-second frame and ending with a 108-second frame.

After reconstruction, the motion corrected time series for each tracer was summed across time to create a single 3D PET emission image. Within individual participants, the sum images for each tracer were aligned to each other using rigid body registration (14). After alignment, the sum images were averaged to create a mean image for each tracer. The mean [$^{18}$F]FDG image was then brought into alignment with the T1-weighted image using rigid body registration and a contrast-preserving gradient vector-field algorithm (15). The final linear transformation was computed by minimizing the error between the forward ([$^{18}$F]FDG → T1) and backward (T1→ [$^{18}$F]FDG) transformations (16). The same procedure was used to align the [$^{15}$O] sum images to the [$^{18}$F]FDG sum image. The computed transformations were combined and used to resample each PET time series in MNI152 2mm atlas space. Nonlinear registration between each T1-weighted image and MNI152 space was performed using the Symmetric Normalization transformation model (17) in ANTS (18).

Relative PET

Standardized uptake values ratios (SUVRs) were computed using specific time windows for each tracer. For [$^{18}$F]FDG, a time window from 40 to 60 minutes post injection was chosen. A sixty second window, starting approximately after the bolus reached the brain, was used for the [$^{15}$O]$H_2O$ and [$^{15}$O]$O_2$ data. A sixty-second window starting 180 seconds after the bolus reached the brain was used for the [$^{15}$O]CO scans. All SUVR images were computed in native space and then resampled to MNI-152 2mm space. To minimize the impact of vascular artifact on our SUVR measurements of oxygen metabolism, a voxelwise spatial regression was run using the resampled [$^{15}$O]$O_2$ SUVR as a dependent variable and [$^{15}$O]$H_2O$ and [$^{15}$O]CO SUVR as independent variables (19). The [$^{15}$O]$O_2$ SUVR was adjusted by subtracting from it the product of the [$^{15}$O]CO SUVR and its regression coefficient. An SUVR approximation of OEF (rOEF) was calculated by dividing the adjusted [$^{15}$O]$O_2$ SUVR by the product of the [$^{15}$O]$H_2O$ SUVR and its regression coefficient. Finally, a SUVR estimate of the relative oxygen-to-glucose index (rOGI) was computed by dividing the [$^{15}$O]$O_2$ SUVR by the [$^{18}$F]FDG SUVR.

Quantitative PET

Voxelwise noise precluded compartmental modeling at the voxel level. Instead, we modified the Hierarchical-Basis Function Method (H-BFM) introduced by Rizzo et al. (20). This model restates the reversible two-compartment model as:

$$C_i(t) = \alpha_1 C_p(t) * e^{-\beta_1 t} + \alpha_2 C_p(t) * e^{-\beta_2 t} + V_b C_a(t) \qquad \text{(Eq. 1)}$$

where $C_i(t)$ is the tissue tracer contraction at time $t$, $C_p$ and $C_a$ are the plasma and whole-blood arterial input functions, and $\alpha_1$, $\alpha_2$, $\beta_1$, and $\beta_2$ are combinations of the model rate constants. If the values for $\beta_1$ and $\beta_2$ are known, the other parameters can be solved using standard linear least squares. In the H-BFM model, the values for $\beta_1$, and $\beta_2$ are determined using a regional constraint. First, the PET image is segmented into several large ROIs and non-linear least squares is used to fit Eq. 1 to the mean time-activity curve for each region. Next, for every voxel within that region, 20 possible $\beta_1$ and $\beta_2$ values are determined by drawing from a normal distribution whose mean and standard deviation are the regional estimate of $\beta_1$ or $\beta_2$ and their standard error. At each voxel, linear least squares is then used to fit Eq. 1 for every possible combination of $\beta_1$ and $\beta_2$. The combination that gives the smallest least squared error is then used to determine $\alpha_1$, $\alpha_2$, $\beta_1$, $\beta_2$, and $V_b$.

One limitation of the H-BFM is that only $\beta_1$ and $\beta_2$ are spatially constrained. Therefore, we modified H-BFM to include a ridge regression constraint:

$$\hat{b} = argmin_b.\{r^T r + \lambda(b - b_{roi})^T (b - b_{roi})\} \qquad \text{(Eq. 2)}$$

where $r$ is the model residuals using equation Eq 1, and $b$ and $b_{roi}$ are vectors containing the linear parameters $\alpha_1$, $\alpha_2$, and $V_b$ for the current voxel and its parent region. The amount of regularization is determined by the value of $\lambda$. After testing a range of $\lambda$ values, we set it equal to the mean counts across the whole brain. For voxels where the estimated CMRglc was 3 median absolute deviations from the median, we reran the fit using a $\lambda$ of 10 times the mean whole brain counts. To account for the difference in scale between $\alpha_1$, $\alpha_2$, and $V_b$, we divided $b - b_{roi}$ by the standard error for each parameter in $b_{roi}$ before computing Eq.2.

Model fitting using non-weighted nonlinear least squares implemented in SciPy (21). For the purposes of the H-BFM model, we used FSL's FAST (22) to segment each visit's mean [$^{18}$F]FDG image into three ROIs whose voxels had similar intensities. Prior to model fitting, the time delay between the arterial samples and the [$^{18}$F]FDG data was accounted for by fitting a one-compartment model to the first sixty seconds (post bolus arrival) of the [$^{18}$F]FDG data (23).

Arterial input function

The arterial input function during the [$^{18}$F]FDG scan was obtained by hand drawing samples through the arterial line. Initial samples were drawn at 2-3 second intervals for at least 120 seconds; all draw and measurement times were recorded with one-second precision. In a smaller subset of participants, an automatic blood sampler (Twilite; Swisstrace) was used to measure arterial input functions for the $^{15}$O PET scans. However, the number of participants with $^{15}$O input functions was very limited; therefore no quantitative $^{15}$O PET data are reported.

Lumped constant modeling

Because [$^{18}$F]FDG is an analog of glucose, a correction factor, called the lumped constant (LC), must be used to convert CMRglc measured with [$^{18}$F]FDG to what would be measured if glucose itself was used as the tracer. As the LC decreases only slightly during hyperglycemia (1), we chose to use the same value (0.81) for both conditions. However, to explore what impact a change in the LC could have our results, we conducted a supplementary analysis where the lumped constant was adjusted using the method of van Golen et al. (24):

$$LC = -0.0043 \cdot C_p + 0.8315 \qquad \text{(Eq. 3)}$$

where in this case $C_p$ is the plasma glucose concentration in mmol·L$^{-1}$. Note that we modified the van Golen et al. method so that the LC at euglycemia ($C_p$ = 90 mg·dL$^{-1}$) was equal to 0.81.
